# Multi-modal mass spectrometry imaging reveals single-cell metabolic states in mammalian liver

**DOI:** 10.1101/2022.09.26.508878

**Authors:** Hua Tian, Presha Rajbhandari, Jay Tarolli, Aubrianna M. Decker, Taruna V. Neelakantan, Tina Angerer, Fereshteh Zandkarimi, Jacob Daniels, Helen Remotti, Gilles Frache, Nicholas Winograd, Brent R. Stockwell

**Affiliations:** Department of Chemistry, Pennsylvania State University, University Park, PA 16802, US; Department of Biological Sciences, Columbia University, New York, NY 10023, US; Ionpath Inc., Menlo Park, CA 94025, US; Department of Chemistry, Columbia University, New York, NY 10027, US; The Luxembourg Institute of Science and Technology, L-4362 Esch-sur-Alzette, Luxembourg; Department of Pharmaceutical Biosciences, Uppsala University, SE-751 05 Uppsala, Sweden; Department of Pathology, Columbia University, New York, NY 10032, US

**Keywords:** Single-cell spatial omics, Mass spectrometry imaging (MSI), Water cluster ion beam secondary ion mass spectrometry imaging ((H_2_O)_n>28K_-GCIB-SIMS)), Desorption electrospray ionization (DESI), Metabolic states, Liver tissue, Molecular, cellular heterogeneity

## Abstract

We have developed a powerful workflow to imaging endogenous metabolism in single cells on frozen tissue, allowing us to discover new cell subtypes in human liver. Performing spatially integrated multiomics in single cells within tissues is at the leading frontier in biology but has been prevented by technological challenges. We developed a critical new technology, cryogenic water cluster ion beam secondary ion mass spectrometry imaging ((H_2_O)_n>28K_-GCIB-SIMS)) at 1 µm single-cell resolution. This allowed us to perform multi-modal mass spectrometry imaging (MSI) to detect metabolites, lipids, and proteins in single cells within functional liver zones and diverse cell types in the native tissue state. Our workflow utilizes the desorption electrospray ionization (DESI) mass spectrometry imaging (MSI) to build a reference map of metabolic heterogeneity and zonation across liver functional units. Then cryogenic (H_2_O)_n>28K_-GCIB-SIMS and C_60_-SIMS integrated metabolomics, lipidomic and proteomics, - characterizing the metabolic state in single cells on the same tissue section. We found for the first time that lipids and metabolites can classify liver metabolic zones and liver cell types beyond histological and protein-marker annotation. This provides a multi-modal workflow to define single-cell states in normal physiology and disease in mammalian tissue.

## Introduction

Recent developments in spatial omics technologies have improved how cellular heterogeneity and tissue organization can be assessed. Efforts such as Human Biomolecular Atlas Program (HuBMAP) (Hu, 2019), Human Tumor Atlas Network (HTAN) (Rozenblatt-Rosen et al., 2020) and Kidney Precision Medicine Project (KPMP(El-Achkar et al., 2021)) aim to map the diversity of cell types and subtypes and their biomarkers to characterize cell types within anatomical structures in healthy and disease tissues at single cell resolution (Borner et al., 2021). While spatial transcriptomics and proteomics have been the most common technologies implemented for such efforts, spatial metabolomics could illuminate the crucial role of effector metabolites that reflect the network of upstream molecules at genomic, transcriptomic and proteomic levels, but have been difficult to detect (Eberlin, 2014; Hickey et al., 2022; Radtke et al., 2020). Mass spectrometry imaging (MSI) is the primary method for spatial metabolomics, enabling multiplexed mapping of untargeted and targeted molecular networks within cells and tissue. Widely used MSI technologies include desorption electrospray ionization (DESI) (Takats et al., 2005) and matrix-assisted laser desorption ionization (MALDI) (Caprioli et al., 1997; Winograd, 2018). DESI is an ambient ionization technique with preferential ionization and characterization of metabolites and lipids at spatial resolution as high as 30 µm (Eberlin, 2014). MALDI is utilized for lipids, abundant proteins, and metabolites with a suitable matrix at an achievable spatial resolution of 5-10 µm (Djambazova et al., 2020).

However, the practical application of MSI at a single-cell resolution has not been possible. First, detection limits hinder imaging of low concentration biomolecules as the amount of material sampled is decreased exponentially within smaller pixels. Second and critically, cryogenic analysis of native cell states is not widely applicable to different MSI tools, which is essential to preserve the pristine chemical gradients in tissue, especially for dynamic and transient metabolites. Finally, due to the incompatibility of sample preparation and difficulty in preserving dynamic metabolic gradients (Tian et al., 2021), it is challenging to acquire multi-omic data within the same tissue section, or to spatially co-localize molecules at single cell level.

We have solved these long-standing problems by developing a buncher-time-of-flight (ToF) SIMS coupled with high voltage water cluster ion beam, (H_2_O)_n(n>28K)_-GCIB and establishing a dual-SIMS workflow. The first step takes advantage of intact biomolecular imaging (up to m/z 5000) at an achievable spatial resolution of ∼1 µm, using a newly designed GCIB operating at ∼70 kV (Sheraz et al., 2019; Tian et al., 2021; Tian et al., 2019). Along with cryogenic sample handling, frozen-hydrated biosamples are transferred in a contamination-free nitrogen atmosphere and images at near-nature status (at 100 K). Cell-type-specific and tissue structure proteins are mapped in the same sample stained with a panel of lanthanides-conjugated antibodies using C_60_-SIMS at the spatial resolution of 1 µm. This development allowed us for the first time to simultaneously map multiple metabolites, lipids, and peptides in the same tissue section at single-cell resolution, with high sensitivity (Taylor et al., 2021).

We applied our pipeline to liver tissue, which is a metabolic hub performing diverse and critical functions, including uptake and storage of nutrients, metabolism, bile secretion, detoxification, protein synthesis, and immune functions (Trefts et al., 2017). To perform these functions, the hepatic parenchyma exhibits metabolic zonation based on the gradient of oxygen-and-nutrient-rich blood along the portal triad (PT) to central vein (CV) axis. The periportal (PP) region surrounding the PT receives a maximum amount of nutrients and oxygen, and is a key site for oxidative metabolism, including β-oxidation, gluconeogenesis, bile formation, and cholesterol synthesis. The pericentral (PC) region surrounding the CV receives less oxygen and nutrients, and is a key site for detoxification, ketogenesis, lipogenesis, glycolysis, glycogen synthesis and glutamine synthesis(Gebhardt and Matz-Soja, 2014; Kietzmann, 2017).

Although histologically hepatocytes in different zones appear homogeneous using conventional imaging methods, these regions are metabolically distinct, shown by the zonal preferences of metabolic enzymes involved in oxidative energy, carbohydrate, lipid, and nitrogen metabolism, including carbamoylphosphate synthetase (CPS1), glutamine synthetase (GS) and cytochrome P450, family 2, subfamily AE, polypeptide1 (Cyp2ae1) (Lamers et al., 1989). The heterogeneity of the liver extends to the cellular level, where multiple cell types are organized in a structured manner to carry out essential liver functions. Within the liver parenchyma, hepatocytes make up 70% of the cell population and are metabolically heterogeneous along the porto-central axis. Non-parenchymal cells, including macrophages (Kupffer cells), liver sinusoidal endothelial cells, and hepatic stellate cells, are located within the hepatic sinusoids and play a critical role in interacting with and regulating parenchymal cells(Aizarani et al., 2019). While liver zonation and function has mainly been explored with immunohistochemistry and single cell spatial transcriptomics(Gebhardt and Matz-Soja, 2014; Halpern et al., 2017), measurement of lipids and metabolites, the effector molecules of metabolic pathways, is lacking.

Here, we present a multimodal MSI approach centered on DESI, (H_2_O)_n_-GCIB-SIMS and C_60_-SIMS for mapping of untargeted, endogenous, unlabeled metabolites and lipids, and targeted protein markers within anatomical structure and cell types, providing a powerful new understanding of metabolic zonation and cell type identity in mouse and human liver. This workflow is complemented and validated by histologic (hematoxylin and eosin (H&E)) staining for anatomical structure annotation within liver and multimodal image alignment, RNAScope for mRNA transcript profiling of landmark genes (Wang et al., 2012) for zonation marking and antibody validation, and MALDI Orbitrap MSI. Moreover, ultra-high performance liquid chromatography (UPLC) coupled with electrospray ionization (ESI) tandem mass spectrometry (MS/MS) is adapted for lipid confirmation in DESI and SIMS experiments. Heterogeneities in liver tissue were detected by the combination of metabolites, lipids, and proteins in human and mouse liver, including sex-specific zonation patterns, functional-specific zonation, and metabolic states of different cell types. We found for the first time that lipid and metabolite composition classify liver zonation and cell types, as well as novel cell subtypes that have not previously been detected by protein or RNA markers. Thus, this new technology for single-cell multi-omic imaging allows for unprecedented understanding metabolic cell states and cell identity.

## Results and discussion

### Multimodal imaging centered on novel mass spectrometry imaging technologies reveals a spatial multiomic (metabolite, lipid and protein) atlas of liver tissue

To comprehensively assess the spatial organization of liver tissue with regard to its cellular and metabolic and lipidomic heterogeneity, we performed multi-modal imaging on consecutive tissue sections of mouse and human liver using DESI, H&E staining, SIMS, and RNAscope, followed by computational data processing for cellular and structure segmentation, omics integration and discriminant analysis. Histological features of the H&E tissue sections were analyzed to guide MSI, with a focus on the PT and CV regions.

DESI was employed to image an entire tissue section at a spatial resolution of 40 µm in both positive and negative ion modes. More than 100 lipid and metabolite features were extracted, and ion species localized to the PP and PC regions (Figure 1a) were identified. A region containing both PT and CV was selected on consecutive tissue sections for single-cell resolution SIMS imaging in a frozen-hydrated state (Figure 1B). We implemented a novel technological development using a three-step process: first, cryogenic (H_2_O)_n_-GCIB-SIMS was performed to localize >100 lipid and metabolites in a pristine native environment at 3 µm per pixel, followed by staining with a panel of lanthanide-labeled antibodies specific to liver biology (Table S1 and S2) on the same tissue section. C_60_-SIMS was then utilized to image metal-labeled antibodies to map cell-types, cell states, and structure-specific protein markers at 1 µm per pixel. As (H_2_O)_n_-GCIB-SIMS removes about 100 nm material from the tissue surface, the antibody markers stain the same cells as those from which lipids and metabolites are detected. This unique approach facilitated image alignment and cell segmentation to register untargeted metabolites and lipids and targeted proteins to single cells on the same tissue section, elucidating biomolecular complexity within different cell types directly in their tissue content without dissociation, which is otherwise not possible in single-cell multimodal image integration.

**Figure 1.**
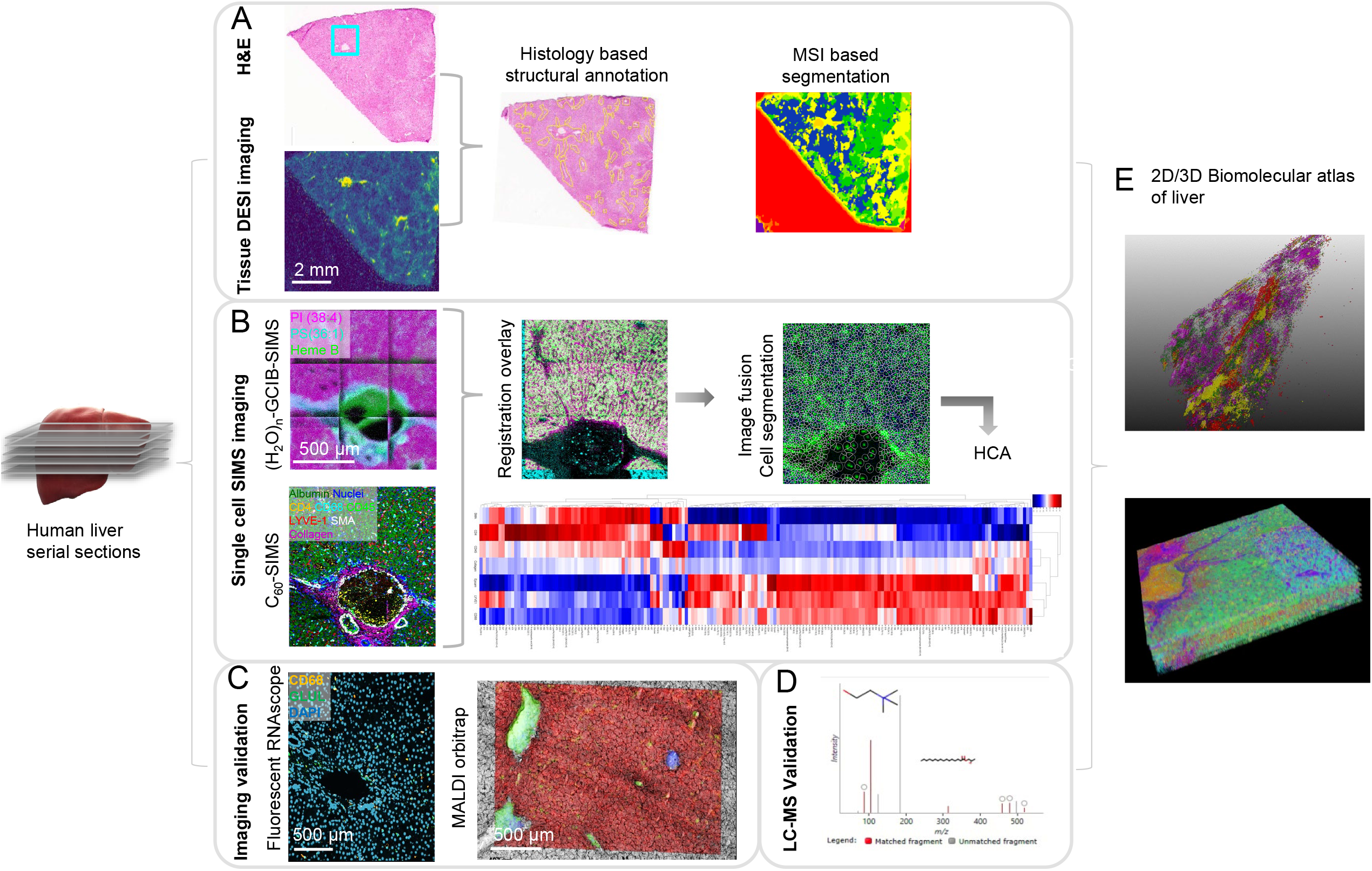
Schematics of MSI-centric multimodal imaging workflow reveals 2D/3D biomolecular atlas of human liver. Consecutive sections from liver tissue blocks are assessed by **A**, DESI-MSI exhibiting the distribution of lipids and metabolites within histologically defined structural units of liver within the tissue architecture and **B**, SIMS-MSI including (H_2_O)_n_-GCIB-SIMS for lipid and metabolite imaging at single cell resolution followed by C_60_-SIMS on the same tissue section, followed by image integration and single cell specific lipid and metabolite extraction **C**, Validation of lipids detected by DESI and SIMS and **D**, structural elucidation by UPLC-MS_E_ lipidomics on liver tissue section homogenate **E**, The 2D images from DESI and SIMS are aligned for 3D reconstruction and visualization.

To validate DESI and SIMS imaging in terms of assignment of zones and cell types, RNAScope was performed on a consecutive tissue section for transcriptomic analysis of several zonation and cell-type-specific markers. To validate ion species assignment by accurate mass and fragmentation pattern, we performed (a) MALDI on an Orbitrap, performed on the serial sections for in situ MS/MS analysis, and (b) lipidomics analysis using UPLC with ion mobility time-of-flight MS^E^ (HDMS^E^) data-independent acquisition and analysis (Figure 1C).

With 2D/3D image reconstruction, molecular and cellular heterogeneity was visualized (Figure 1E), showing distinct molecular clusters in different areas and Correlation of proteins and metabolites and lipids in the PT region in human liver.

### DESI MSI reveals metabolic zonation-specific metabolites and lipids in mouse liver

To evaluate metabolite and lipid distribution with metabolic zones, we first examined histology of mouse liver tissue sections obtained from six different mice (3 male and 3 female). H&E-stained images were annotated to identify the PT and CV regions, showing uniform distribution patterns within the tissue (Figure 2 Ai, S1a). The annotation was then validated by transcript and protein markers targeting metabolic enzymes that are specific to PP and PC regions(Lamers et al., 1989). For example, Albumin (*Alb*) was concentrated in the PP region, and Glutamine Synthetase (*Glul*) was localized in the PC region using RNAScope, respectively (Figure 2 Aii, S1b). The distributions were consistent with the previous work(Halpern et al., 2017), however, we noted sex-linked differences. *Alb* was concentrated more around the PP zone in male mice, whereas it expanded towards the central vein in female mice. *Glul* expression has been previously shown to exhibit sex-specific variation in different mouse and rat strain(Sirma et al., 1996); we observed a similar distribution pattern between male and female mice in the C57BL6 mouse strain used in this study.

**Figure 2.**
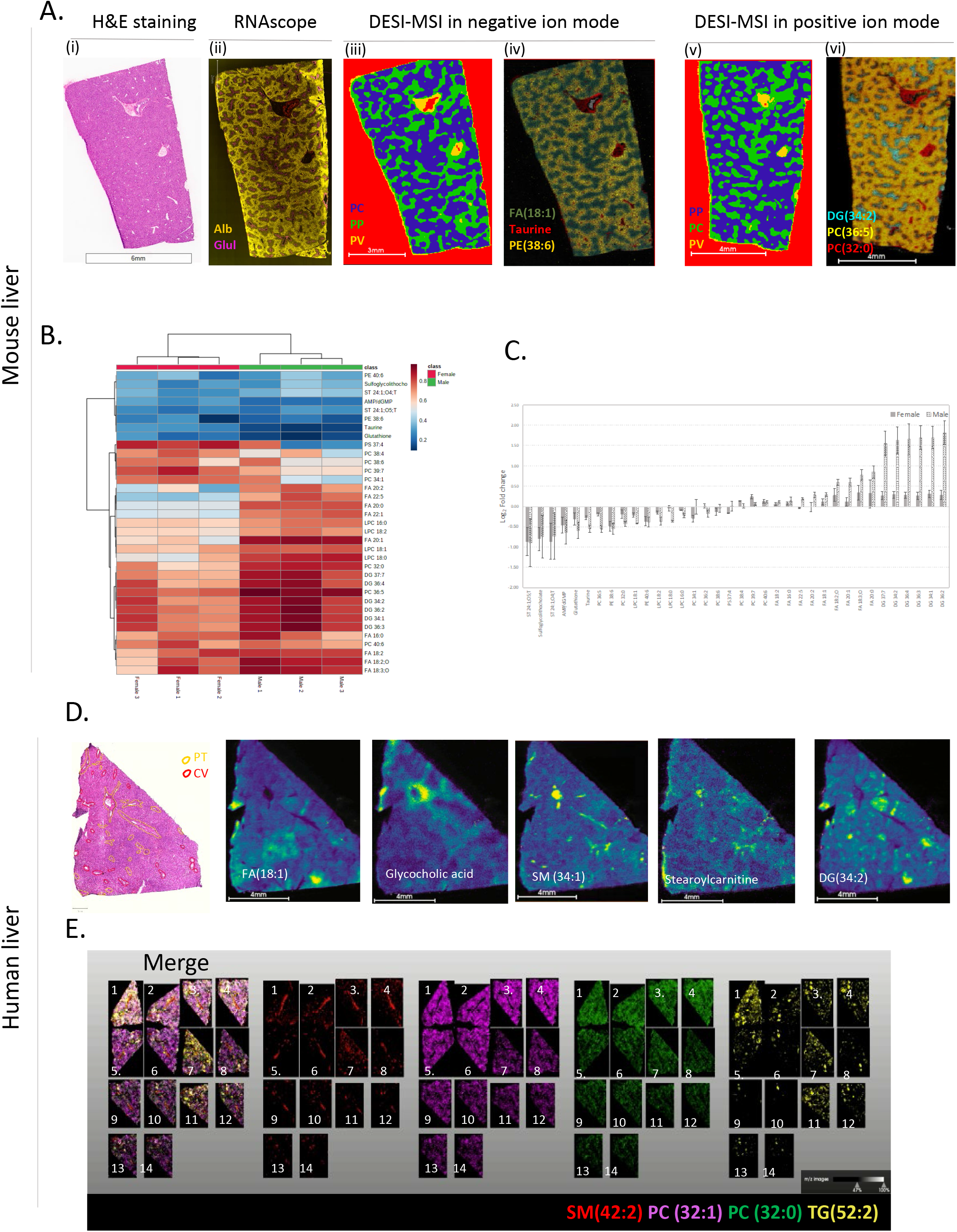
DESI MSI reveals periportal and pericentral specific lipids and metabolites in mouse and human Liver:. **Ai**, H&E staining performed on a normal mouse liver section to identify the central vein (CV) and portal triad (PT) regions. **Aii**, RNAScope is performed on a consecutive tissue section stained for expression of Albumin (Alb-yellow) and Glutamine Synthetase (Glul-magenta) showing differential staining for periportal (PP) and pericentral (PC) regions respectively. **Aiii, Av**, Spatial segmentation of pixels based on distribution of a few lipids and metabolites from DESI-MSI in (Aiii) negative and (Av) positive ion mode. (Aiv, Avi) Distribution of a few lipids and bolites showing PC and PP specificity in both ion modes are shown. **B**, Heatmap depicting the predictive performance of features measured by AUC-ROC (in rows) and classification (in columns) from DESI for PP and PC regions in female (n=3) and male (n=3) mice liver sections. **C**, Fold change values comparing the mean intensity of PC versus PP regions for the features from (B) Error bars represent mean ± standard deviation. **D**, H&E section with manual annotation of CV and PT regions in human liver section. The corresponding images for distribution of different classes of lipids are shown. **E**, Distribution of lipids in consecutive sections for 3-dimensional visualization of lipids as shown in Supp video1.

DESI imaging of mouse liver showed distinct mass spectra representing metabolites and lipids in both positive and negative ion mode. In positive mode, ions corresponding to lipids from different classes, including glycerophospholipids and glycerolipids were observed (Figure S2a, Table S3). In negative ion mode, free fatty acids and small metabolites in a low m/z range (m/z 100-400), glycerophospholipids in high m/z range (m/z 700-800) and conjugated bile acids were observed (Figure S2b, Table S3). In order to extract PP and PC specific lipids and metabolites, we first sought to create a segmentation mask for these regions within tissue sections. A few ion species (*m/z* 124.01, 280.23, 282.25, 306.07 and 309.28) in negative ionization mode and *m/z* 534.29, 631.47, 734.57, 794.51, 802.5 and 852.55 in positive ionization mode) exhibiting unique distribution patterns were used to perform bisecting k-means clustering with Euclidean distance, within the SCiLS software platform. Spatial segmentation was performed with edge-preserving denoising to remove pixel-to-pixel variability observed in MSI datasets(Alexandrov et al., 2010). This resulted in PP and PC clusters in individual mouse liver tissue sections in both positive and negative ion mode, shown as a segmentation map (Figure 2 Aiii and Av, S1c and S1d).

We confirmed cluster specificity by comparing *Alb* and *Glul* transcript distribution with DESI segmentation (Figure S1b, S1c and S1d). To identify lipids and metabolites that discriminate PC versus PP clusters, we implemented a binary classifier—receiver operating characteristic (ROC)-area under the curve (AUC) curve analysis(Bradley, 1997; Mandrekar, 2010). Each tissue-specific ion was evaluated for its discriminating power, using a threshold of the area under the curve (AUC) >0.70 to be considered a classifier. We observed that metabolites such as glutathione (GSH), taurine, and conjugated bile acids were zonated in PP region (Figure 2B, S3c, S3d). Our results were complementary to previous results on the metabolic gradients from PC to PP. Liver bile acids are primarily synthesized in PC hepatocytes, they flow towards the bile duct and the enzymes cascade involved in bile acid biosynthesis is organized spatially. In particular, the enzyme conjugating bile acids is abundant in PP region, where taurine also comes in from blood supply(Halpern et al., 2017; Ikeda et al., 2012).

As noted, we observed sexual dimorphism in the distribution of several lipid species. While previous studies have shown sex-specific differences in lipid metabolism, it has not been shown in the spatial context within liver (Mittendorfer, 2005; Soares et al., 2017). Diacylglycerols (DGs) and free fatty acids (FAs) are zonated in the PC region (Figure 2B, S3a, S3b), with the average ROC-AUC for individual DG ions for males of >0.89 and females >0.7 (Figure 2B). Compared to females, male mice showed higher amounts of DGs in PC versus PP zones (log_2_ fold change of ∼1.5 for males and 0.4 for females) (Figure 2C). Similar observations were made for FAs (Figure 2B, 2C, S3b). While some FAs such as FA(18:1), FA(20:1), FA(18:3;O), FA(18:2) and FA(20:0) show high specificity for PC versus PP zone, with the average ROC-AUC for individual ions for males of >0.88 and females >0.7 (Fig 2B), others such as FA(18:2;O), FA(16:0), FA(22:5) and FA(20:2) exhibited higher specificity for PC zones in males versus females, with the average ROC-AUC for individual ions for males of >0.73 and females <0.57. FA (20:0) and FA(18:3;O) showed relatively higher abundance in PC versus PP in both male and female mice, and FA(20:1) and FA(22:5) showed minimal differences between the regions in female mice (Figure 2C, S3b).

The PC region has previously been reported to exhibit a higher degree of lipogenesis, fatty acid synthesis and acetyl CoA carboxylase expression than the PP zone(Guzman and Castro, 1989; Quistorff et al., 1992). On the other hand, phospholipids, including phosphatidylcholine (PCs) and lysophosphatidylcholine (LPCs), did not exhibit a specific pattern. Some lipids (e.g., PC(40:6)) showed similar distribution between the PP and PC zones, while others (e.g., PC (36:5), PC (32:0), PC(34:1)) exhibited differential distributions between regions, or between regions and sexes (e.g., LPC(18:2), LPC(18:0), LPC(16:0), PC(38:4) and PC(39:7)) (Figure 2B, 2C, S3e). Thus, while male and female mice have histologically and morphologically similar livers, they are metabolically quite distinct.

### DESI MSI reveals zonation-specific metabolites and lipids in human liver

The established DESI workflow was then performed on consecutive tissue sections from human liver tissue. H&E staining was performed to assess tissue morphology and annotate PP and PC regions (Figure 2D). RNAScope was used to validate *Glul* staining, which unlike in mouse liver, appeared as dispersed puncta around the CV region (Figure S4). This is likely due to lower expression of the gene in human liver. Hence, PP and PC regions were annotated manually based on H&E staining (Figure 2D).

DESI imaging at 40 µm spatial resolution showed similar average mass spectra profiles in both positive and negative ion modes compared to mouse liver (Figure S2c, S2d). Taurine-and-glycine-conjugated bile acids were abundant in the PT and PP region, as well as in the septa connecting the neighboring PTs, showing similar functional zonation between human and mouse liver (Figure 2D, S5b). The vasculature were identified by the abundance of the marker, Heme B. Cholesterol, a precursor to bile acids, was concentrated within PV and CV (Figure S5e). Sphingomyelins (SMs) including SM 34:1 were colocalized with cholesterol (Figure 2D, S5e), where SMs are postulated to form hydrophobic lipid raft domains with cholesterol, preventing hepatic damage from bile salts, and also playing a role in pathophysiology(Amigo et al., 1999; Simons and Ehehalt, 2002). Similarly, stearoylcarnitine was localized in the PP region (Figure 2D), where acylcarnitines play an important role in transferring long-chain fatty acids to mitochondria for β-oxidation(Longo et al., 2016). Several small metabolites and different lipid species exhibited differential distributions in human tissue (Figure 2D, S5). FAs, DAGs and triglycerides (TAGs) exhibited distinct distribution patterns, which unlike in mouse tissue, did not overlay well with the PC region (Figure 2D, S5a, S5c, S5d). This could stem from the complexity of human liver, where metabolic gradients are more dynamic and heterogeneous, based on a person’s genetic profile and changes in gene expression and metabolic-enzymes-based factors such as diet, hormones, gender and underlying pathology (Geisler and Renquist, 2017; Kietzmann, 2017). Similarly, varied patterns of phospholipid species were observed within the liver (Figure S5f).

In addition to 2D imaging of tissue sections, 3D DESI aids the visualization of the distribution of lipids and metabolites within three-dimensional tissue structures and gradients with respect to tissue depth, capturing branching of blood flow and anatomic structure. The resulting 3D model visualizing differentially distributed lipids is shown in Supplementary Video 1.

### SIMS imaging delineates heterogeneities of multiple biomolecules and cell types at single-cell level in mouse liver

To associate lipids and metabolites with different types of cells in the liver, dual SIMS imaging at the resolution of 1-3 µm was performed on a consecutive mouse liver sections. Guided by H&E stained images, a region of interest (ROI) containing both CV and PV was selected (Figure 3 Ai) for (H_2_O)_n(n=30k)_-GCIB-SIMS imaging in negative ion mode, mapping more than 100 lipids and metabolites in both mouse and human liver. The highly heterogenous metabolites and lipids were observed around CV and PV, including nucleotides, bile acids, glucose, fatty acids (FA), phosphatidylinositols (PIs), phosphatidylserines (PSs), lysophosphatidylserines (LPSs), phosphatidic acids (PAs), lysophosphatidic acids (LPAs). PI(38:4) was highly concentrated in the PC region, and PS(40:6) distributed in complimentary locations with an elevated concentration around the PP region. Heme B was mainly present inside of the veins, while taurocholic acid was present in the PP region (Figure 3Aii, S6A and S5E). On the same region, ten cell-type-specific and tissue structure markers illuminated a heterogeneous cell landscape and recapitulated the anatomical structures in the mouse liver (Figure 3 Aii). The single ion images of selected species are detailed in Figure S6, exhibiting various distribution patterns.

**Figure 3.**
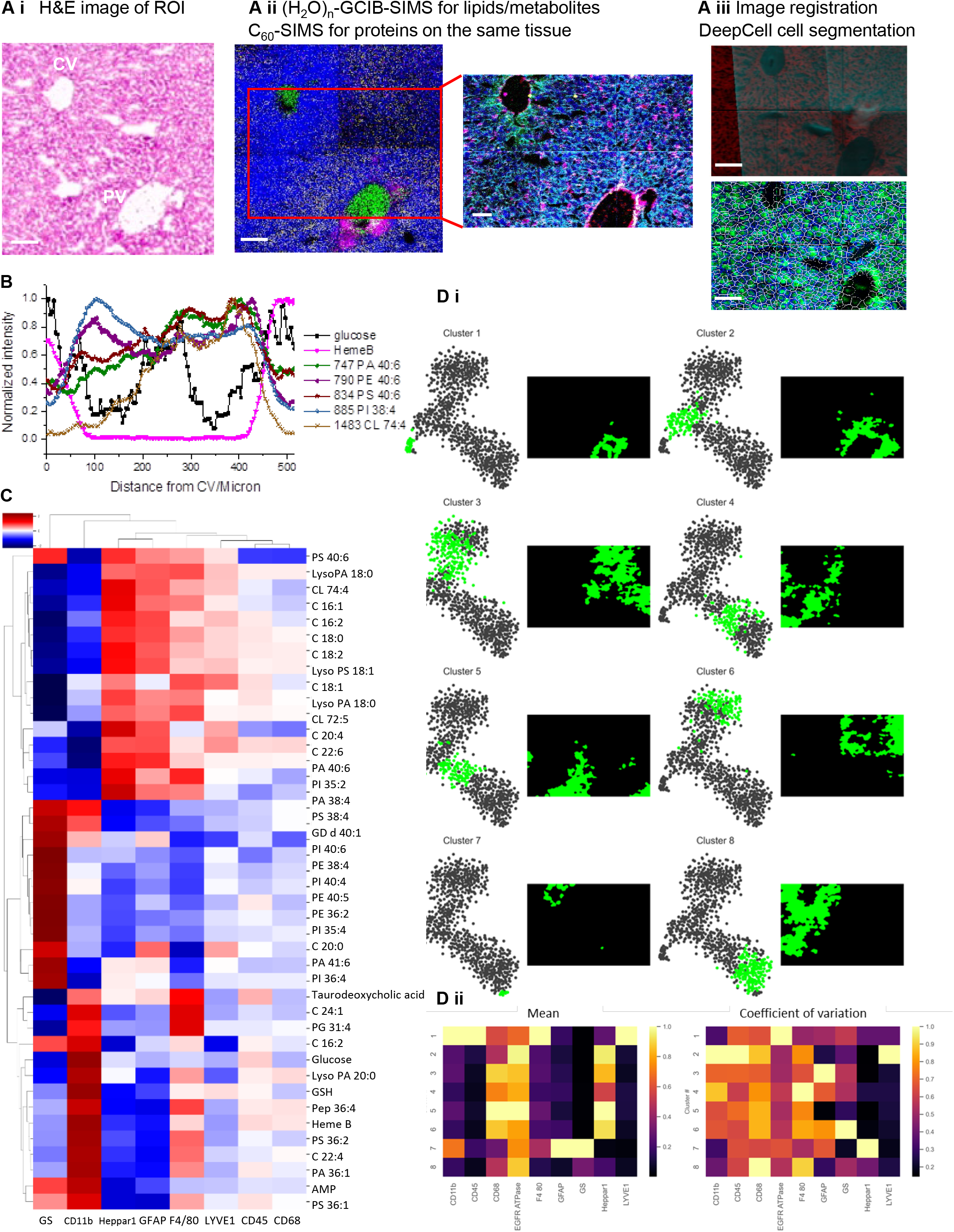
Dual high-resolution SIMS imaging delineates the metabolomic and lipidomic states in different cell types on the mouse liver section. **Ai**, H&E staining image of central vein (CV) and portal vein (PV) region. **Aii**, Representative color overlay images of Dual-SIMS. Metabolites and lipids image in the same region as in Ai on a serial frozen-hydrated section using (H_2_O)_n(n=30k)_-GCIB-SIMS at the spatial resolution of 3 µm, Blue, PI 38:4; Magenta, Taurocholic acid ; Yellow, PS 40:6; Green, HEME B. More single specie images are in Figure S6. Proteins image by lanthanides-conjugated antibodies from the region highlighted in the red box using C_60_-SIMS at the spatial resolution of 1 µm, Red, CD45 and CD11b; Yellow, GFAP; White, LYVE-1; Magenta, F4/80 and CD68; Cyan, EGFR and Na/KATPase; Green, Glul; Blue, Nuclei. Scale bar 100 µm. **Aiii**, Registration of dual-SIMS images for alignment; Single cell segmentation using DeepCell. More details are described in supplementary Figures S16 and S17. **B**, Intensities changes of various species from the center of CV to PV (as the blue line in A i). **C**, HCA map shows the variation of different metabolites/lipids in various types of cells. **Di**, tSNE clustering to classify the cell types as in cluster 1-8 by lipids and metabolites only. **Dii**, Correlation of cluster 1-8 to 9 protein markers.

Principle component analysis (PCA) revealed clusters of ions contributing to the major features around the CV and PV (Figure S7). Heme B, a marker for vasculature, was highly concentrated inside the vein in principal component 2 (PC2). As an exclusive and essential lipid constituent of mitochondrial membranes, a variety of cardiolipins (CLs) was elevated around the PV rather than the CV, which is in line with the oxygen gradient captured in PC4 (Paradies et al., 2014). PC5 highlights the CV region with species PI(38:4), PS(38:4), PA(38:4), LPA(18:0), FA(20:4) and PS(38:3). On the other hand, PC6 shows the radiated gradient around the PV with dominated species LPS(16:1), and LPS(16:0), echoing an earlier study on increased PS-PLA1 or LPS receptor 1 (LPS1) mRNA in hepatocellular carcinoma (HCC) tissues(Uranbileg et al., 2020). The chemical gradients of these species were measured along the porto-central axis by line scanning (Figure 3B). Glucose appeared more abundant inside the CV and PV, the same as Heme B. GSH and AMP had similar patterns as glucose, indicating high energy consumption in the CV and PV region. Most PI species were concentrated around the CV region (e.g., PI(38:4)), while PA and PS species were found to be higher in the PV region (e.g., PA(40:6), PS(40:6)). PE(40:6) appeared to be more intense at the edge of the CV and PV, contributing to the curved structure and mechanical resistance facilitated by the conical shape of PE). Figure S8 demonstrates more species with similar chemical gradients across the CV and PV.

Sequential C_60_-SIMS imaging on the same region profiled by (H_2_O)_n(n=30k)_-GCIB-SIMS further revealed the distribution of targeted protein markers using a panel of lanthanide-labeled antibodies. The panel was designed to identify major cell types, immune cells, cell boundaries and zonation markers within liver (Table S1). Leukocytes, Ito stellate cells, sinusoidal endothelial cells and macrophages/Kupffer cells were localized with the markers CD45/CD11b red, GFAP in yellow, LYVE-1 in white, CD68 and F4/80 in magenta, respectively (Figure 3Aii). As expected, GS (in Green) expression was abundant around CV, which is consistent with RNAScope results (Figure S2b). EGFR and Na/K-ATPase in cyan defined the cell border, facilitating cell segmentation and image alignment to register multiple lipids and metabolites in individual cells. The single-channel imaging of each antibody using C_60_-SIMS (Figure S10) was validated by immunohistochemistry (IHC) (Figure S9).

Image processing for alignment, registration, and cell segmentation by Deepcell(Keren et al., 2018) are described in Figure 3 Aiii and detailed in Figure S15-17. With this, the detected omics molecules using dual-SIMS imaging were registered to individual segmented cells for further statistical analysis. The hierarchical clustering algorithm (HCA) compares the intensities variation of monitored metabolites/lipids among different cell types, providing a comprehensive view of cell signatures and metabolic states. As shown in Figure 3C, GS expressing cells comprise significant PI(38:4), which metabolizes to downstream signaling molecules phosphatidylinositol phosphate (PIPs) known to bind many proteins and control protein-protein interactions(Amos et al., 2019). PE(40:6) also contributes to the GS-positve cells, particularly cells in the inner circle of the CV and PV. Along with PA(38:4), PS(38:4), GD(d40:1), PI(40:6), PE(38:4), PI(40:4), PE(36:2), FA(20:0), PA(41:6) and PI(36:4), these lipids were more abundant in Glul-positive cells around the CV. The glucose content was highest among the CD11b positive cells, suggesting a high glucose-dependant metabolism. Sinusoidal cells labeled by LYVE-1 are rich in lipids with a low degree of unsaturation. Cells expressing the periportal hepatocyte-specific marker (Heppar1/CPS1) consist of a higher level of LPAs and cardiolipins.

Next, t-distributed stochastic neighbor embedding (t-SNE) was performed to cluster the single cell data points that integrate the omics molecules detected using SIMS, showing metabolites and lipids only can classify the cell populations. As in Figure 3D, Clusters 1-8 highlight the regions around the PP and PC regions, particularly cluster 1, identifying the portal endothelial cells that are consistent with the localization of LYVE-1 positive cells (Figure 3Di, 3Dii). The cells in clusters 2-6 are co-localized with Heppar-1 positive cells (Figure 3 Dii). Separating the same type of cells into distinct clusters is likely related to the vastly different metabolic/lipidomic states of hepatocytes at various locations, and reflects the varying functions among the same cell types. Cluster 7-8 are around the CV, the exact location of Glul expressing cells (Figure 3 Di, 3 Dii). Hence, the abundance of metabolites and lipids allows for clustering of cells into distinct subpopulations, some of which are not captured by conventional protein markers.

### SIMS delineates heterogeneity by cell-type specific multi-omics at single cell level in human liver

The same workflow for dual SIMS imaging was repeated around PT on a human liver tissue section (Figure 4). The high-resolution images demonstrated the heterogeneous biomolecules, as PS (36:1) outlined the PT, PI (38:4) was localized outside the PT primarily, and Heme B was inside of the PT (Figure 4 Aii). The single ion images of 246 metabolites and lipids are in Figure S11, among them, 120 species were annotated.

**Figure 4.**
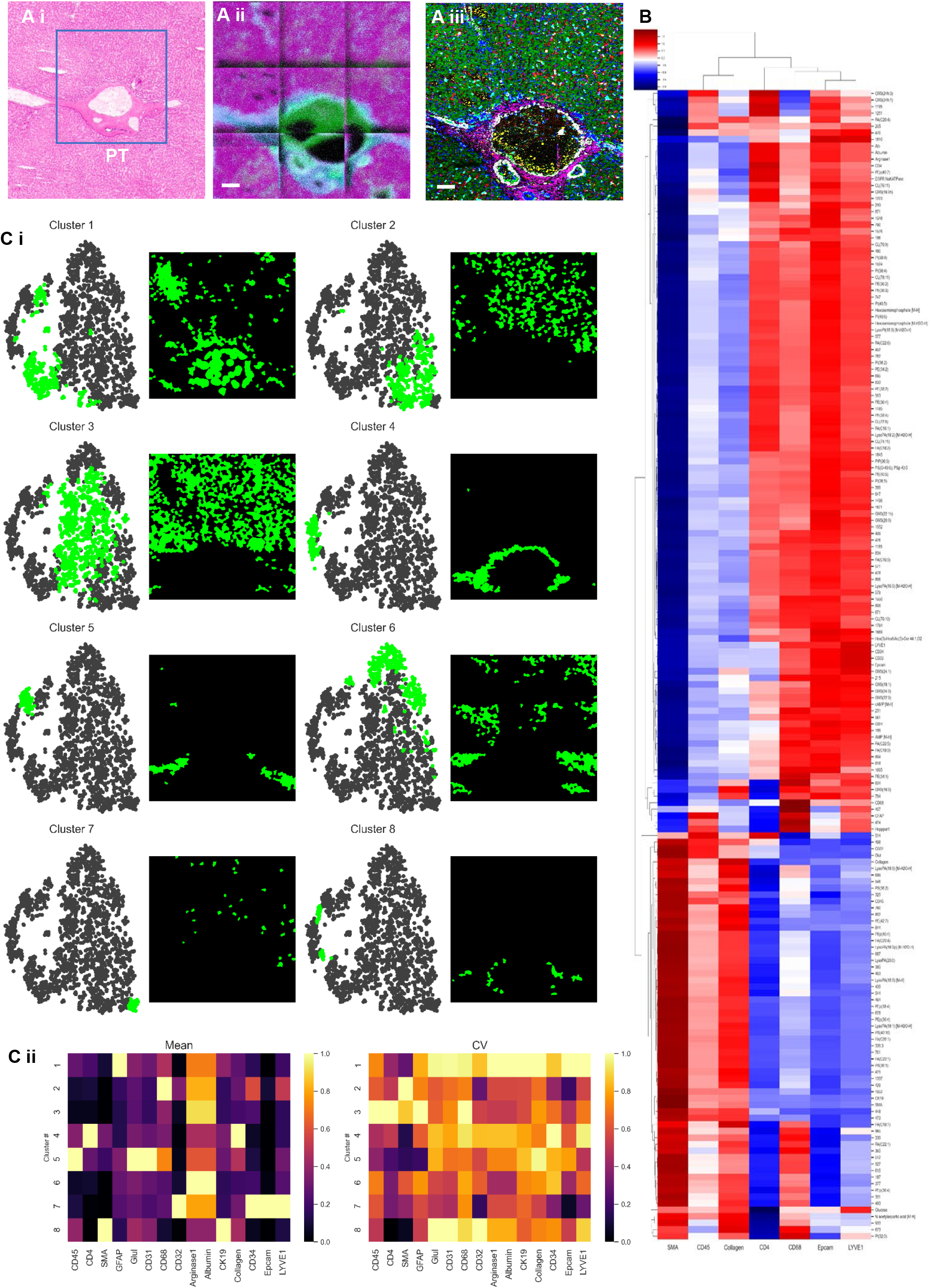
Dual high-resolution SIMS imaging delineates the metabolomic and lipidomic states in different cell types on the human liver tissue section. **Ai**, H&E image. The selected region contains the central vein and portal vein. **Aii**, Representative color overlay images of Dual-SIMS. Metabolites and lipids image in the same region as in Ai on a serial frozen-hydrated section using (H_2_O)_n(n=30k)_-GCIB-SIMS at the spatial resolution of 3 µm, Blue, PS 36:1; Magenta, PI 38:4; Green, HEME B. More single specie images are in Figure S10. Proteins image by lanthanides-conjugated antibodies from the same region after (H_2_O)_n(n=30k)_-GCIB-SIMS using C_60_-SIMS at spatial resolution of 1 µm, Red, CD45 and CD11b; Magenta, GFAP; White, LYVE-1; Yellow, Heppar-1; Cyan, EGFR and Na/KATPase; Green, Glul; Blue, Nuclei. **Aiii**, The image alignment and registration of dual-SIMS images (i.e., Aii and Aiii). More details are described in supplementary Figures S16 and S17. **B**, HCA map shows the variation of different metabolites/lipids in different types of cells. **Ci**, tSNE clustering to classify the cell types as in cluster 1-8 by lipids and metabolites only. **Cii**, Correlation of cluster 1-8 to 15 protein markers.

PCA analysis distinguished two anatomical features and metabolic flux around the PT (Figure S12). The PT ring has abundant PS(36:1), PA(36:1) and PI(38:4), with significant higher PS(36:1) that defines the PT region. Inside the PT ring, Heme B and several PA species are dominant. The metabolic flux also shows anti-correlated patterns, with taurodeoxycholic acid/taurodeoxycholic, FA (C18:2) and FA (C18:1), taurocholic acid and PEp(40:6) to the left of the PT ring and GSH to the right.

On the same tissue, protein markers show the major cell types, immune cells and cell states (Figure 4 Aiii). With the same computational process (Figure 3 Aiii), HCA elucidates the variation of monitored metabolites/lipids in different cell types and structural regions, namely CD4, CD45, Epcam, LYVE-1, CD68, SMA and collagen I (Figure 4B). IHC validation and single-channel image of antibody panel (Table S2) are detailed in Figure S13 and S14, respectively. T-helper cells (CD4) and leukocytes (CD45), were located mainly around the PT, sharing similar metabolism of high glucose consumption. Epcam, as an epithelial antigen, shows the highest GSH level that correlates to the activation of antioxidant mechanism. Sinusoidal cells (LYVE-1) comprise the higher ganglioside GM3 and PE species, which are highly relevant to cell adherence and mechanical resistance to highly packed cellular region (Calzada et al., 2016; Labrada et al., 2018). Without distinct lipidomic and metabolic features, macrophage/Kupffer cells (CD68) seemed to have slightly higher CLs, representing a higher number of mitochondria. As a major tissue structure marker, SMA is co-localized with PS 36:1 abundant region.

tSNE clustering was further applied to classify cells using metabolites and lipids (Figure 4 Ci and 4 Cii). Cluster 1, 4, 5 and 8 were consistent with the GFAP, collagen I, CD45 and SMA positive cells, respectively. Low expression of Arg-1, CD4, Glul and CK19 were also observed in these clusters. Periportal hepatocytes expressing Arg-1 and Albumin were sub-classified into clusters 2, 3, 6 and 7. The overlapping of protein markers for each tSNE cluster was likely to be the shared signature metabolites and lipids among them. Several factors contributed to the success of the cell classification by metabolites and lipids, the successive high-resolution SIMS images on the same tissue that facilitates the accurate cell segmentation and image registration, and cryogenic SIMS workflow that preserves the pristine chemistry in single cells.

### RNAScope, MALDI Orbitrap and LC-MS to validate the core MSI workflow

To validate antibody markers used in the study, namely CD68, Glul, Albumin and LYVE1, single-molecule mRNA fluorescent *in situ* hybridization was performed on human liver tissue section. Color overlaid images of RNAScope show the distribution of the RNA copies (Figure S4) consistent with images of proteins (Figure S9, S10, S13 and S14). For example, protein GS and its corresponding mRNA transcript, *Glul*, were localized around CV, but not PV. In addition, fluorescent slide scanning allowed easy identification of PV and CV, compared to H&E staining in mouse liver tissue sample (Figure S1b).

The identity of the zone-specific lipids was assessed using accurate mass match of the MSI-derived ions of interest against the library databases(Fahy et al., 2009) followed by UPLC-MS^E^ fragment analysis from lipidomics analysis from liver tissue lipid extracts (Table 1). The analysis of DESI and lipidomics analysis with lockmass correction helps to align the precursor mass with high accuracy. The UPLC-MS^E^ data was searched for the most commonly formed adducts, [M+ H^+^, Na^+^, K^+^]^+^ in positive ion mode for DESI and [M-H]^−^ for negative ion mode for both DESI and SIMS. The metabolites were annotated by accurate mass search against the library database(Smith et al., 2005; Wishart et al., 2022) and accurate mass and ion mobility drift time of the standard when available. Most of the precursor ion matches between the DESI and LC-MS were phospholipids and glycerolipids in positive ion mode, and for SIMS were phospholipids in negative ion mode. Similarly, confirmation of lipid species based on MALDI is shown in Table S3 and S4. DESI and SIMS identified a complementary list of species around CV and PV (Table 1), resulting from the difference in preferential ionization for different lipid species. High-resolution SIMS uncovered several metabolites and lipids that have a thin circulate structure around both central and portal veins, such as GSH, AMP, PE(40:6), PA(36:1), PS(38:3) and PI(38:4); some species were much more concentrated inside the veins, such as LPAs (Figure S6 and Figure S11). While DESI confirmed distinct distributions of more free fatty acids, conjugated bile acids and glycerolipids (Figure 2 and Figure S3). All species were validated by in situ tandem MS using MALDI (Table S3 and S4). However, metabolites and fatty acids are not validated by MALDI due to the mass interferences for species below *m/z* 350 from the matrix. The results demonstrate the complementary approach using a variety of MSI methods for more comprehensive imaging of biosamples.

**Table 1.**
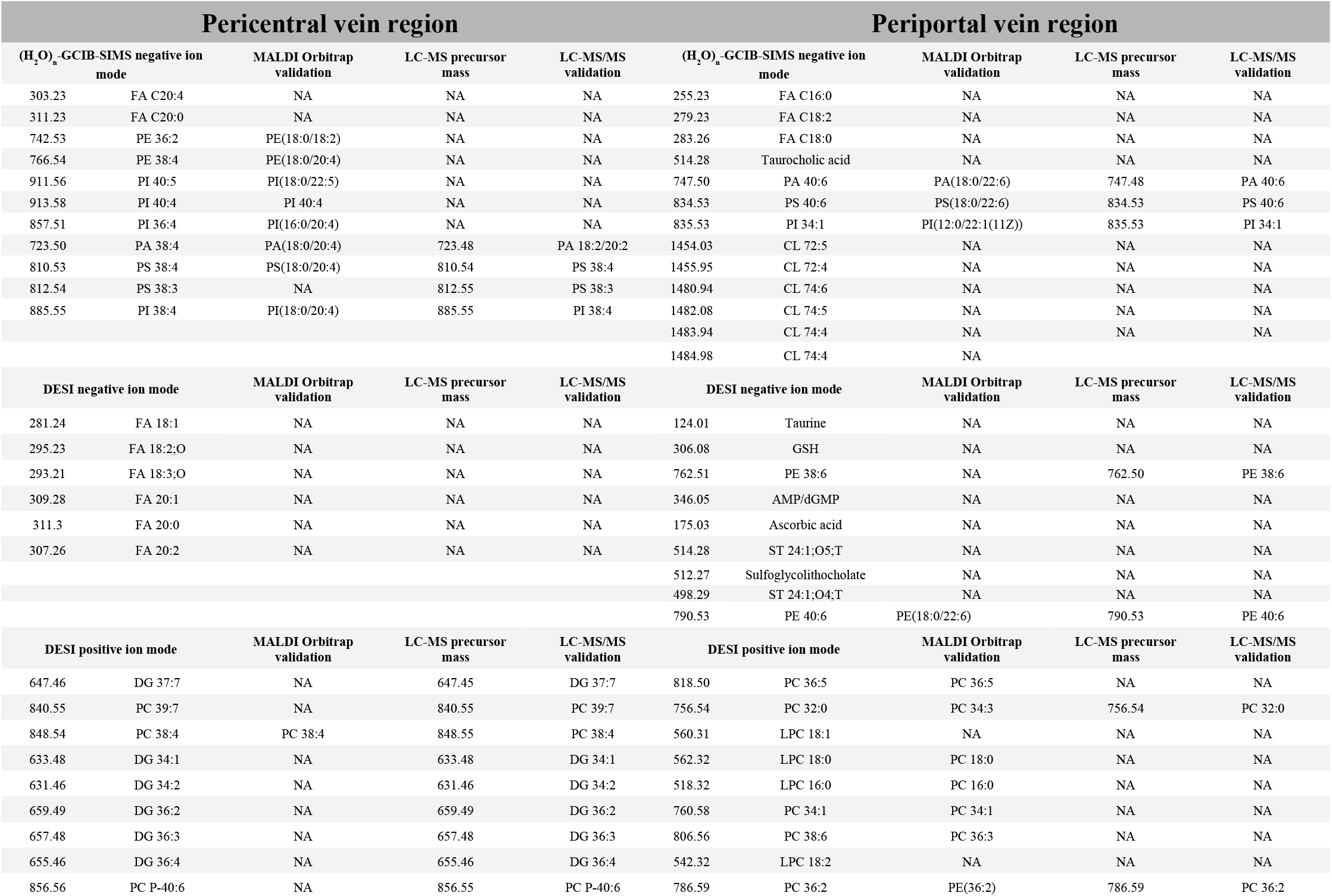
Validation of lipid ion species from SIMS and DESI-MSI with MALDI-MSI and LC-MS/MS.

| Pericentral vein region |  |  |  |  | Periportal vein region |  |  |  |  |
| --- | --- | --- | --- | --- | --- | --- | --- | --- | --- |
| (H<sub>2</sub>O)<sub>n</sub>-GCIB-SIMS negative ion mode |  | MALDI Orbitrap validation | LC-MS precursor mass | LC-MS/MS validation | (H<sub>2</sub>O)<sub>n</sub>-GCIB-SIMS negative ion mode |  | MALDI Orbitrap validation | LC-MS precursor mass | LC-MS/MS validation |
| 303.23 | FA C20:4 | NA | NA | NA | 255.23 | FA C16:0 | NA | NA | NA |
| 311.23 | FA C20:0 | NA | NA | NA | 279.23 | FA C18:2 | NA | NA | NA |
| 742.53 | PE 36:2 | PE(18:0/18:2) | NA | NA | 283.26 | FA C18:0 | NA | NA | NA |
| 766.54 | PE 38:4 | PE(18:0/20:4) | NA | NA | 514.28 | Taurocholic acid | NA | NA | NA |
| 911.56 | PI 40:5 | PI(18:0/22:5) | NA | NA | 747.50 | PA 40:6 | PA(18:0/22:6) | 747.48 | PA 40:6 |
| 913.58 | PI 40:4 | PI 40:4 | NA | NA | 834.53 | PS 40:6 | PS(18:0/22:6) | 834.53 | PS 40:6 |
| 857.51 | PI 36:4 | PI(16:0/20:4) | NA | NA | 835.53 | PI 34:1 | PI(12:0/22:1(11Z)) | 835.53 | PI 34:1 |
| 723.50 | PA 38:4 | PA(18:0/20:4) | 723.48 | PA 18:2/20:2 | 1454.03 | CL 72:5 | NA | NA | NA |
| 810.53 | PS 38:4 | PS(18:0/20:4) | 810.54 | PS 38:4 | 1455.95 | CL 72:4 | NA | NA | NA |
| 812.54 | PS 38:3 | NA | 812.55 | PS 38:3 | 1480.94 | CL 74:6 | NA | NA | NA |
| 885.55 | PI 38:4 | PI(18:0/20:4) | 885.55 | PI 38:4 | 1482.08 | CL 74:5 | NA | NA | NA |
|  |  |  |  |  | 1483.94 | CL 74:4 | NA | NA | NA |
|  |  |  |  |  | 1484.98 | CL 74:4 | NA |  |  |
| DESI negative ion mode |  | MALDI Orbitrap validation | LC-MS precursor mass | LC-MS/MS validation | DESI negative ion mode |  | MALDI Orbitrap validation | LC-MS precursor mass | LC-MS/MS validation |
| 281.24 | FA 18:1 | NA | NA | NA | 124.01 | Taurine | NA | NA | NA |
| 295.23 | FA 18:2;O | NA | NA | NA | 306.08 | GSH | NA | NA | NA |
| 293.21 | FA 18:3;O | NA | NA | NA | 762.51 | PE 38:6 | NA | 762.50 | PE 38:6 |
| 309.28 | FA 20:1 | NA | NA | NA | 346.05 | AMP/dGMP | NA | NA | NA |
| 311.3 | FA 20:0 | NA | NA | NA | 175.03 | Ascorbic acid | NA | NA | NA |
| 307.26 | FA 20:2 | NA | NA | NA | 514.28 | ST 24:1;O5;T | NA | NA | NA |
|  |  |  |  |  | 512.27 | Sulfoglycolithocholate | NA | NA | NA |
|  |  |  |  |  | 498.29 | ST 24:1;O4;T | NA | NA | NA |
|  |  |  |  |  | 790.53 | PE 40:6 | PE(18:0/22:6) | 790.53 | PE 40:6 |
| DESI positive ion mode |  | MALDI Orbitrap validation | LC-MS precursor mass | LC-MS/MS validation | DESI positive ion mode |  | MALDI Orbitrap validation | LC-MS precursor mass | LC-MS/MS validation |
| 647.46 | DG 37:7 | NA | 647.45 | DG 37:7 | 818.50 | PC 36:5 | PC 36:5 | NA | NA |
| 840.55 | PC 39:7 | NA | 840.55 | PC 39:7 | 756.54 | PC 32:0 | PC 34:3 | 756.54 | PC 32:0 |
| 848.54 | PC 38:4 | PC 38:4 | 848.55 | PC 38:4 | 560.31 | LPC 18:1 | NA | NA | NA |
| 633.48 | DG 34:1 | NA | 633.48 | DG 34:1 | 562.32 | LPC 18:0 | PC 18:0 | NA | NA |
| 631.46 | DG 34:2 | NA | 631.46 | DG 34:2 | 518.32 | LPC 16:0 | PC 16:0 | NA | NA |
| 659.49 | DG 36:2 | NA | 659.49 | DG 36:2 | 760.58 | PC 34:1 | PC 34:1 | NA | NA |
| 657.48 | DG 36:3 | NA | 657.48 | DG 36:3 | 806.56 | PC 38:6 | PC 36:3 | NA | NA |
| 655.46 | DG 36:4 | NA | 655.46 | DG 36:4 | 542.32 | LPC 18:2 | NA | NA | NA |
| 856.56 | PC P-40:6 | NA | 856.55 | PC P-40:6 | 786.59 | PC 36:2 | PE(36:2) | 786.59 | PC 36:2 |

## Discussion

This study demonstrates for the first time major technological improvements that enable a multi-modal workflow for multiplexed imaging of metabolites, lipids and proteins for integrated spatial omics at anatomical structure and single-cell resolution. The multi-scale biomolecule detection works efficiently to distinguish anatomical features (*e*.*g*., CV, PV and PT) and metabolic zones in liver tissue. Moreover, significant variations of species are observed in different types of liver cells (*e*.*g*., hepatocytes, Kupffer, sinusoidal cells, macrophages, T cells and leucocytes), demonstrating that different metabolic states are needed for spatial division of labor to efficiently manage a multitude of metabolic functions. Finally, we observed sex-specific differences in the distributions of many lipid species, suggesting that male and female livers may be functionally quite distinct, with implications for understanding sex-specific differences in disease risk.

There is growing interest in dissecting cellular and metabolic heterogeneity directly on tissue at single-cell resolution. Spatial metabolomics is becoming an attractive approach to answer questions about liver metabolic reprogramming, which is associated with a spectrum of liver diseases. This requires characterizing cell types and cell neighborhoods, as well as an atlas of biomolecule abundances within the cell types, taking into account where these cell types exist within the tissue architecture. However, there are few analytical tools to image multiple types of molecules in single cells directly on tissue without dissociation. Consecutive SIMS imaging on the same frozen-hydrated tissue offers single-cell resolution and high chemical sensitivity to integrate spatial multi-omics (untargeted metabolomics and lipidomics and targeted proteomics) in the same individual cells and at a near-native state of biological tissue. With well-preserved gradients of small molecules, which would be otherwise diffused by any chemical fixation and drying process, the metabolic state of different types of cells can be revealed in liver tissue. In addition, cell population separation on tissue directly by metabolic states has not been reported, but some success has been shown on classification of co-cultured cells by metabolic states using MALDI imaging and computational approach SpaceM(Rappez et al., 2021).

With cryogenic multi-modal SIMS imaging at high-resolution and data processing, we show for the first time that integrated metabolomic and lipidomic profiling in individual cells can be used for cell classifications without protein markers and tissue dissociation. Additionally, subsets of cell types are captured by distinct metabolic signatures among cells expressing the same proteins. As the role of individual lipid species are understudied, this platform could be extended to overlay spatial transcriptomics and proteomics characterizing the distribution of enzymes involved in lipid metabolism to fully elucidate the spatial diversity and its functional significance. This will provide a rare opportunity to investigate previously unknown cellular subtypes and their unique protein-lipid-metabolite interactions.

A robust pipeline with high-resolution multi-modal MSI and computational processing adds layers of spatial omics about metabolic and lipidomic heterogeneity in different types of cell populations, distinguishing subpopulations and distinct metabolic functions within individual cells on normal liver tissue. This workflow can be readily applied to liver disease models for the discovery of metabolic vulnerability and associated cell types, heterogenous shift of disease microenvironment, and cell-to-cell interaction, ultimately leading to new therapeutic opportunities.

## Acknowledgments

This research was supported by NIH HubMAP (1UG3CA256962-01). We thank our colleague, Dr. Andrea J. Radtke from NIH who provided insight and expertise in liver antibody panel. We would also like to show our gratitude to Dr. Dan Graham from the University of Washington and Dr. Nathan H. Patterson from Vanderbilt University for sharing their expertise in MSI data processing.

## Author contributions

HT, PR and BRS designed the multimodal imaging workflow for liver tissue study. HT conceived the dual-SIMS imaging workflow, established the lanthanide-conjugated antibody panel and immunostaining protocol, and performed tissue immunostaining, SIMS data acquisition and general/discriminant processing. PR optimized sample processing for multimodal imaging, performed sample preparation, DESI imaging and data analysis, lipidomic experiment and data analysis and multimodal data comparison. JT developed the software platform and algorithm for data processing, image registration and segmentation under the supervision of HT. AMD performed RNAScope imaging and analysis. TVN performed cryosectioning and assessed IHC data with PR. TA performed MALDI imaging. FZ contributed to lipidomic experiments. TVN and JD performed mouse tissue sample preparation. HR performed tissue identification, pathological analysis and annotation of tissue structure. NW provided the SIMS instrumentation. HT, PR and BRS drafted and edited the paper. BRS supervised the project. HT and PR contributed equally.

## STAR Methods

### RESOURCE AVAILABILITY

#### Lead contact

Further information and requests for resources and reagents should be directed to and will be fulfilled by the lead contact, Hua Tian.

#### Materials availability

This study did not generate new unique reagents.

#### Data and code availability

##### Data availability

Mass spectrometry imaging date are in standardized imzML format. SIMS data is available at doi:10.26207/6a38-tr35. DESI and SIMS data will also be available from https://portal.hubmapconsortium.org by end of July 2022. The software license is provided for analyzing the data using Ionoptika Analyser for up to 7 days. Please contact Ionoptika if you wish to use the software beyond 7 days via our Support e-mail. Accession numbers are listed in the key resources table.

- Any additional information required to reanalyze the data reported in this paper is available from the lead contact upon request.

## METHOD DETAILS

### Tissue preparation

Normal human liver sample from a 51-year-old female was retrieved from the Columbia University tissue bank, under a protocol approved by the Institutional Review Board and stored at -80°C until use. Mouse liver tissues were derived from 17 weeks old male (n=3) and female (n=3) C57/BL6 mice. The excised liver tissues were flash frozen on dry ice filled with hexane, and stored at - 80°C until use. Small blocks of both human and mouse tissue were cryosectioned at 8-10µm thickness and thaw mounted on microscope glass slides for H&E, DESI and RNAScope analysis and gold-coated slides for SIMS analysis. For multimodal imaging, consecutive sections for DESI, H&E, SIMS and RNAScope were placed on the respective slides and stored at -80°C until analysis.

### Histology

Tissue sections were stained with hematoxylin and eosin (H&E) staining at Columbia University Molecular Pathology Shared Resource facility, scanned at 20X magnification and histological examination was performed by a pathologist to annotate the anatomical structures.

### DESI MSI data acquisition and analysis

All the experiments were performed on Synapt G2-Si QToF mass spectrometer (Waters, Milford, MA), coupled to a DESI ion source. Data was acquired in sensitivity mode in both positive and negative ion mode. The DESI parameters used were capillary voltage and sampling cone voltage of 0.65kV and 50V respectively, scan time of 0.145 sec/pixel, pixel size of 40 µm^2^, DESI sprayer angle of 75°, nebulizing gas (N_2_) pressure of 0.3 PSLM. The solvent used was methanol: water 95:5 (v/v) with 0.01% formic acid and 20pg/µl leucine enkephalin, at a flow rate of 1.5 µl/min. Tissue sections were dried in a desiccator for ∼10 min prior to analysis. Peak picking and lockmass correction using the protonated ion of leucine enkephalin ([M+H]^+^, m/z 556.2771) or the deprotonated molecular ion ([M-H]^−^, m/z 554.2615) was implemented in MassLynx software (Waters, version 4.1). The centroided data files were converted to mzml using msconvert from Proteowizard(Kessner et al., 2008) followed by conversion to imzml format using imzMLConverter(Race et al., 2012). The imzml files were imported into the SCiLS lab software (Bruker, version 2021c) and subsequent data analysis was performed. Total ion count (TIC) normalization was performed and up to n peaks were selected of m/z intervals of ±0.03Da were selected. Spatial segmentation analysis was performed using bisecting k-means clustering on edge-preserving denoised data. Area under the receiver operator characteristic curve was also performed within the SCiLS platform.

### Untargeted Lipidomics sample preparation

20 mg of liver was homogenized using beadrupter and lipids were extracted using 1050ul of 1:2 ratio of ice-cold methanol containing 0.01% w/v butylated hydroxyl toluene and dichloromethane, vortex mixing and incubating the samples overnight at -20C. One volume of ice-cold water was added and mixed for phase separation followed by centrifugation. The lower organic phase containing lipids were collected in a new vial, dried under a gentle stream of nitrogen and stored at -80C until analysis. The samples were reconstituted in isopropanol: acetonitrile: water at the ratio of 11:9:2 v/v/v before analysis.

### Chromatographic separation and mass spectrometry analysis

Lipidomics experiments were performed on Synapt G2-Si mass spectrometer equipped with Acquity UPLC system (Waters, Milford, MA) in both positive and negative electrospray ion modes. The chromatographic separation was performed on Acquity UPLC BEH300 C18 column (1.7um particle size, 2.1×100mm) (Waters, Milford, MA) over an 18 min gradient. The column temperature was set at 55C. A binary mobile phase consisted of (A) 60:40 v/v acetonitrile: water and (B) 85:10:5 v/v/v isopropanol/acetonitirile/water, each containing 10mM ammonium acetate and 0.1% acetic acid. The gradient was initiated at 40% B, followed by linear gradient to 50% by 2min ramped up to 99%B by 18min, and the column was equilibrated for 2min to the initial condition. The flow rate was set at 400ul/min and injection volume was 2ul in positive mode and 5ul in negative mode. Data was acquired on high-definition data independent mode with ion mobility (HDMS^E^), over the mass range of m/z 50 to 1200 Da and scan rate of 0.1sec per scan. The parameters used for mass spectrometry data acquisition is as follows: for positive and negative mode, capillary voltage and sampling cone voltage of 2.8kV and 35V; and 2.5kV and 32V were used respectively. The source and desolvation temperature were 120°C and 500°C respectively. Desolvation gas (N_2_) flow was set at 850 L/hr. MS data was calibrated using leucine encephalin infusion at a flow rate of 10µl/min. Default ion mobility settings were used. The low collision energy was set at 4 eV and high energy was 25-60 eV. Mass calibration was performed using sodium formate and collision cross section (CCS) calibration was performed using CCS Major mix (Waters).

### Lipid and metabolite identification

The assignment of lipids and metabolites specific of selected ions from DESI-MSI were based on its assignment based on accurate mass and isotopic pattern score using Progenesis software (Waters Inc., Milford, MA). The accurate mass search against the available databases including Lipidmaps^35^, HMDB(Wishart et al., 2022) and Metlin(Smith et al., 2005) for [M+ (H/Na/K)]^+^ adducts were searched in positive ion mode and [M-H]^−^ adducts in negative ion mode. For UPLC-HDMS^E^ lipidomics data, the assignment of the lipid features was based on retention time information, accurate mass as well as fragmentation information. Fragmentation match was made in Progenesis software, where fragmentation score and match were assessed. It was also performed in MS^E^ dataviewer (Waters, Milford, MA), where the fragments were confirmed against the lipidmaps structure database (LMSD)(Sud et al., 2007) for the adducts mentioned above with mass tolerance of 5ppm. Finally, the accurate mass of DESI-MSI based ion was matched against the UPLC-HDMS^E^ data for assignment.

### Successive SIMS imaging and data processing

#### Cryogenic (H_2_O)_n_-GCIB-SIMS

Both (H_2_O)_n_-GCIB and C_60_-SIMS were performed on a buncher-ToF instrument, J105 3D Chemical Imager (Ionoptika, Southampton, UK. Abbv. J105). The water cluster ion beam is pulsed through a pulser in the gun column, where the distance to the sample surface is 0.533 m. Beam tuning was assisted with an oscilloscope (Tektronix TDS 2024, USA) with detection by a secondary electron detector (SED). The singly-charged (H_2_O)_n_ cluster size at beam energy of 70 kV with a time of flight (ToF) of 103 µs was calculated using the ToF equation as n = 30,900 (Figure S15). The SED offset was 8 µs. Beam focus was measured by scanning a 1000 mesh grid (Agar Scientific, Essex, UK). The average beam spot sizes were calculated using 20/80 percent of maximum intensities and were 1.60±0.01 µm and 1.16±0.45 µm for 70 keV (H_2_O_30k_ ^+^ and 40 kV C_60+_, respectively (Figure S14). The beam dither was adapted to match the image pixel size. The mass resolution m/Δm was 6875 around *m/z* 100, and 10,000∼12,000 up to *m/z* 2000. The live readout of mass resolution was from the software, Ionoptika SIMS Mainframe during the data acquisition.

The gold coated Si wafer with the frozen-hydrated mouse/human liver tissue section was plunged into liquid nitrogen and inserted to the pre-chilled cold sample stage in J105 instrument and kept at 100 K during GCIB-SIMS imaging. This cryogenic sample handling preserved the frozen-hydrated state thus maintaining the chemical gradients in the tissue section.

Guided by the anatomical features on the semi-serial H&E stained section, an area of interest was selected for SIMS imaging in negative ion mode using a 70 keV (H_2_O)_30k_^+^ beam. The acquisition was in negative ion mode with 256 × 256 pixels using a 2 × 2 tiled image mode for mouse liver tissue sections, or 768×768 pixels using a 3 × 3 tiled image mode for human liver tissue sections. Each tile covers 400 × 400 µm^2^ (3.1 µm per pixel) for each section. With 1 pA of beam current and 296 shots per pixel, the ion doses were 3.01×10^12^ ions/cm^2^ each tile.

#### Immunostaining

The antibody panel, designed to identify the major cell types, immune cells, cell proliferation, structure and nuclei within the liver is described in Table S1 and S2. Briefly, after the (H_2_O)_n_-GCIB-SIMS profiling was performed, the frozen tissue was placed at -20 °C and 4 °C consecutively for 1 h each for temperature equilibration, followed by fixation in 4 % formalin solution at 4 °C for 30 min. Non-specific protein binding was blocked with 3 % BSA (Bovine Serum Albumin) for 45 min at room temperature. Overnight staining was then performed with the antibody cocktail solution (750 ug/mL for each antibody) at 4 °C. The stained slide was washed with 0.2 % Triton X-100 in PBS (phosphate-buffered saline) 1X for 8 min before the final nuclear staining with Intercalator-Ir at 300 µL/section. After washing with double-distilled water for 10 min and air-drying for 30 min, the slide was again inserted into the SIMS instrument, this time for C60 imaging. For immunohistochemistry, briefly, tissue sections were fixed in cold acetone, washed and incubated with 30% hydrogen peroxide. The sections were blocked with 10% goat serum and incubated with primary antibody for 90 mins followed by biotinylated secondary antibody at room temperature, with washing steps in between. This was followed by addition of avidin-biotin complex reagent and then DAB (3,3′-Diaminobenzidine) with washing between steps. The stained slide was washed with water and counterstained with hematoxylin, mounted with coverslip and scanned at 20x resolution.

#### C_60_-SIMS

High resolution images using 40 keV C_60+_ were then acquired on the same area previously profiled by the (H_2_O)_n_-GCIB-SIMS. The acquisition was conducted in positive mode to localize various cell types. This was achieved by spatially detecting unique *m/z* ions of the isotopic metal tags associated with cell-specific antibodies. To resolve single cells, the C_60_^+^ beam was finely focused to 1.0 µm to image roughly the same area which has been analyzed by (H_2_O)_n_-GCIB. With the beam current of 5 pA and 1000 shots per pixel, the ion dose was 8.57×10^13^ ions/cm^2^. The dwell time was 100 ms/pixel. The lanthanide tags from eight antibodies and the nuclear marker were detected at an adequate signal intensity to allow co-registration with lipid and metabolite ions detected by (H_2_O)_n_-GCIB-SIMS.

#### SIMS Data processing

Single mass channels from tiled C_60_ and (H_2_O)_n_-GCIB-SIMS images were extracted using [Ionoptika’s Analyze] and subsequently used for downstream processing, all performed with custom developed Python code. Co-registration of C_60_ and (H_2_O)_n_-GCIB-SIMS images was by first selecting mass channels that demonstrated a representative morphology of the tissue and normalizing each to an intensity range of [0, 1]. Normalized images were registered using SimpleITK(Beare et al., 2018) (v 2.0.2) to determine the best affine transform between the C_60_ (fixed) and H_2_O (moving) images by minimizing the mean square difference using a gradient descent optimizer (Figure S16). All H_2_O channel data was then transformed to the C_60_ image space. The general nuclear and membrane channels from the C_60_ data set were then used to segment single cells using DeepCell(Bannon et al., 2021) (v 0.9.0). Since the (H_2_O)_n_-GCIB-SIMS data has been registered to the C_60_ image space, the segmentation instances can be used to extract integrated counts of species in both SIMS data sets. Integrated protein expression from C_60_-SIMS images was used to determine thresholds for cell classification (marker positive or negative) (Figure S17). Hierarchical clustering analysis (HCA) was performed with seaborn (v 0.11.1) on the integrated lipid and metabolite mass channels from the (H_2_O)_n_-GCIB-SIMS data set, using the cell types determined from the C_60_ image data (Figure S18).

#### MALDI-MSI

Chemicals and solvents (analytical grade) were purchased from the following sources: α-Cyano-4-hydroxycinnamic acid 98% (CHCA) (Sigma Aldrich), 1,5-Diaminonaphthalene 97% (DAN) (Sigma Aldrich), acetonitrile (ACN) (Honeywell), chloroform (Acros Organics), methanol (Carl Roth), and trifluoroacetic acid (TFA) (Sigma Aldrich). All chemicals used in this study were stored, handled, and disposed of according to good laboratory practices (GLP).

Mouse and Human liver sections on ITO glass (Sigma, Milwaukee, WI, US) were stored at -80ºC until analysis. Prior to matrix application, the tissue sections were removed from the freezer, placed on a cold steel plate (-20ºC) and freeze-dried in a desiccator for 30 minutes. The combination of steel plate and desiccator was efficient for removing the water from the tissue without compromising its structural integrity and limiting the migration of analytes. On the mouse liver section, DAN matrix (10 mg/ml, ACN:H_2_O 7:3) was applied. For the analysis of human liver, 2 section were coated in matrix, one with DAN (same as mouse) for negative ion mode and one with CHCA matrix (5mg/ml, CHCl_3_:MeOH 1:1) for positive ion mode analysis. Application was performed with an HTX TM sprayer (HTX Technologies LLC, USA), temperature: 30ºC (Dan)/ 40ºC (CHCA), passes: 8 (DAN)/16 (CHCA), flow rate: 0.12 ml/min, velocity: 1200 mm/min, drying time: 2 s, line spacing 2.5 mm.

AP-MALDI analysis was performed using an AP-MALDI UHR ion source (Masstech Inc., USA), which has been described in detail elsewhere,^40,41^ coupled to an LTQ/Orbitrap Elite high-resolution mass spectrometer (Thermo-Fisher Scientific, USA) in positive and negative ion mode. For imaging, the AP-MALDI source was operated in “Constant Speed Raster” motion mode. To explore the detectable species and instrument settings for both ion modes, one whole mouse liver section was analyzed interlaced in positive and negative ion mode with a laser beam diameter of 20 µm and a stepping size of 50 µm, laser settings 2500 Hz, 5%. Spectrum acquisition parameters were 800 ms maximum injection time, mass range: 500 – 2000 Da and mass resolution: 120k at m/z 400. Human liver was analyzed with higher spatial resolution (15 µm laser spot and stepping size) and positive (CHCA matrix, laser settings 500 Hz, 10%) and negative (DAN matrix, laser settings 1500 Hz, 5%) ion mode analysis was performed on 2 separate, consecutive sections. Spectrum acquisition was adjusted to 500 ms maximum injection time, mass range: 350 – 1550 Da and mass resolution: 120k at m/z 400. Species identification was performed with on-tissue tandem-MS with a 1.5 Da isolation window, and collision-induced dissociation/ higher-energy collision dissociation (CID/HCD) was performed with collision energies of 27-45%, adjusted for each species individually. Tandem-MS scans were summed up over 30-120 seconds. Data analysis and visualization was performed with Thermo Xcalibur 2.2 (Thermo-Fisher Scientific, USA), MultimagingTM (ImaBiotech, France), METASPACE,^42^ and LipostarMSI (Molecular Horizons Srl, Italy).^43^ Lipid identification was performed in LipostarMSI (database: LIPIDMAPS, mass accuracy: 2 ppm; mass and isotopic pattern score: 80%+).

#### RNAScope

We captured transcription distributions of select liver cell marker genes via *in situ* hybridization of specific targeting probes with the RNAscope Multiplex Fluorescent v2 Assay Protocol^27^ optimized for fresh-frozen samples. Our modifications to the commercial protocol included using half-concentration wash buffer (0.5X) for all wash steps downstream of probe incubation, and excluding the recommended protease step entirely. In mouse tissue, we spatially detected the transcripts for, ALB (Albumin), GLUL (Glutamine synthetase), and PTPRC (Protein Tyrosine Phosphatase Receptor Type C), and in human tissue, we spatially detected transcripts for GLUL, CD68 (Macrophage Antigen CD68), and LYVE1 (Lymphatic Vessel Endothelial Hyaluronan Receptor 1).

Tissue sections were first post-fixed with 4% paraformaldehyde (PFA) in phosphate buffered saline (PBS) and dehydrated in Ethanol (EtOH) immediately after fixation, immersed for 5 minutes at a time in 50% EtOH, 70% EtOH, 100% EtOH, and 100% EtOH an additional time. Samples were then air-dried and treated with RNAscope^®^ Hydrogen Peroxide Reagent for ten minutes at 23°C to 25°C and washed twice with deionized water. Importantly, we excluded the commercial protease step because tissue integrity was lost, and we could achieve stronger signal without any protease treatment. These steps constitute the pretreatment steps.

These pretreated sample slides were incubated with prewarmed target probes (20 nmol/L of each oligo probe) overnight. In mouse tissue, ALB was targeted with RNAscope® Probe-Mm-Alb-C2 (ACD;Cat No. Cat No. 437691-C2), GLUL was targeted with RNAscope Probe-Mm-Glul (ACD;Cat No. 426231), and PTPRC was targeted with RNAscope® Probe-Mm-Ptprc-C3 (ACD;Cat No. 318651-C3). In human tissue, GLUL was targeted with RNAscope® Probe-Hs-GLUL-No-XMm (ACD;Cat No. Cat No. 511171), CD68 was targeted with RNAscope® Probe-Hs-CD68-C4 (ACD;Cat No. 560591-C4), and LYVE1 was targeted with RNAscope® Probe-Hs-LYVE1 (ACD;Cat No. 426911).

The tissue was incubated in the primary target probes overnight (18-21 hours) at 40°C inside the HybEZ hybridization oven (ACD). After overnight probe hybridization, samples were incubated in Amplifier 1 (preamplifier) (2 nmol/L) in hybridization buffer B (20% formamide, 5× SSC, 0.3% lithium dodecyl sulfate, 10% dextran sulfate, blocking reagents) for 30 minutes; Amplifier 2 (2 nmol/L) in hybridization buffer B at 40°C for 15 minutes; and Amplifier 3 (label probe) (2 nmol/L) in hybridization buffer C (5× SSC, 0.3% lithium dodecyl sulfate, blocking reagents) for 15 minutes. After each hybridization step, slides were washed with 0.5X wash buffer (0.05× SSC, 0.015% lithium dodecyl sulfate) two times at room temperature. Chromogenic detection was performed utilizing a horseradish peroxidase (HPR) construct specific to each gene-dedicated imaging channel and a fluorescent Opal reagent of choice. For the mouse sections, ALB was stained with Opal 520 Reagent (Perkin Elmer, FP1487001KT), GLUL was stained with Opal 570 Reagent (Perkin Elmer, FP1488001KT), and PTPRC was stained with Opal 690 Reagent (Perkin Elmer, FP1488001KT). For the human sections, GLUL and LYVE1 were both stained with Opal 520 Reagent (Perkin Elmer, FP1487001KT), thus needing to be imaged in separate tissue sections, and CD68 was stained with Opal 570 Reagent (Perkin Elmer, FP1488001KT). Each Opal reagent dye was diluted 1:1500 in RNAscope^®^ Multiplex TSA Buffer. Nuclei were stained with DAPI (4′,6-diamidino-2-phenylindole) and coverslips were mounted over slides in Fluoro-Gel (EMS; 17985-10) and imaged by spinning disc confocal microscopy and Aperio Versa 8 fluorescent slide scanner.

## KEY RESOURCES TABLE

| REAGENT or RESOURCE |  |  | SOURCE | IDENTIFIER |
| --- | --- | --- | --- | --- |
| Antibodies for mouse liver |  |  |  |  |
| Metal tag | Target | Clone/Host | Cell/Pathway | Manufacturer/Catalog No. |
| 89Y | CD45 | 30-F11, Rat IgG2b | Pan leukocyte | Fluidigm/3089005B |
| 142Nd | Heppar-1/CPS1 | HepPar1, mouse monoclonal | Periportal hepatocytes (Zone 1) | Novus/NBP3-08970 |
| 143Nd | GFAP | EPR1034Y, Rabbit monoclonal | Ito Stellate Cells | Abcam/ab218309 |
| 147Sm | GS | ab240193,Rabbit monoclonal | Pericentral hepatocytes (Zone 3) | Abcam/ab240193 |
| 152Sm | F4/80 | CI:A3-1, Mouse IgG2b | Kupffer cells | Abcam/ab6640 |
| 155Gd | CD11b | M1/70, Rat IgG | Leukocytes | Ionpath/ 715503-100 |
| 156Gd | CD68 | FA-11, Mouse IgG2b | Macrophages | Abcam/ ab237968 |
| 174Yb | LYVE1 | Rabbit polyclonal | Sinusoidal endothelial cells | Novus/ NB600-1008 |
| 176Yb | EGFR | EP38Y, Rabbit monoclonal | Cell membrane | Abcam/ ab272293 |
| 176Yb | Na/K ATPase | EP1845Y, Rabbit monoclonal | Cell membrane | Abcam/ab167390 |
| 191 Ir | Nuclear DNA |  | Nuclei | Fluidigm/ 201192B |
| Antibody for Human liver |  |  |  |  |
| Metal tag | Target | Clone/Host | Cell/Pathway | Manufacturer/Catalog No. |
| 89Y | CD45 | D9M8I, Rabbit IgG | Pan leukocyte | Cell Signaling/13917S |
| 113 In | CD4 | RPA-T4, mouse monoclonal | T cells and macrophages | Novus/NBP2-25199 |
| 141Pr | SMA | 1A4, Mouse IgG2a | Smooth muscle cells in vascular walls, sinusoildal endothelial cells and activated fibroblasts | Fluidigm/3141017D |
| 143Nd | GFAP | EPR1034Y, Rabbit monoclonal | Ito Stellate Cells | Abcam/ab218309 |
| 145Nd | Heppar-1/CPS1 | HepPar1, mouse monoclonal | Periportal hepatocytes (Zone 1) | Novus/NBP3-08970 |
| 147Sm | GS | ab240193,Rabbit monoclonal | Pericentral hepatocytes (Zone 3) | Abcam/ab240193 |
| 148Nd | CD31 | JC/70A, Mouse monoclonal | Endothelial cells | Abcam/ab264090 |
| 151Eu | CD68 | D4B9C,Rabbit IgG | Macrophages | Cell Signaling/76437S |
| 153Eu | CD32 | FUN-2, Mouse IgG2b | B cells, monocytes, granulocytes and platelets | Fluidigm/3153018B |
| 158Gd | Arginase1 | D4E3M™,Rabbit IgG | Zone 1-2 hepatocytes, and macrophages | Cell Signaling/93668S |
| 161Dy | Albumin | EPR20195, Rabbit monoclonal | Periportal hepatocytes (Zone 1) | Abcam/Ab271979 |
| 166Er | CK19 | SPM561, Mouse monoclonal | Cholangiocytes (Portal triad) | Abcam/ab212569 |
| 169Tm | CD34 | QBEND/10, Mouse / IgG1 | Endothelial cells | Abcam/ ab198395 |
| 170 Er | EpCAM | E6V8Y, Rabbit IgG | Hepatic stem/progenitor cells | Cell Signaling /93790S |
| 171Yb | LYVE1 | EPR21857, Rabbit monoclonal | Sinusoidal endothelial cells and subsets of macrophages | Abcam/ab232935 |
| 176Yb | EGFR | EP38Y, Rabbit monoclonal | Cell membrane | Abcam/ ab272293 |
| 176Yb | Na/K ATPase | D4Y7E, Rabbit IgG | Cell membrane | Cell Signaling/ 23565S |
| 191 Ir | Nuclear DNA |  | Nuclei | Fluidigm/ 201192B |
| 196Pt | Collagen I | EPR7785, Rabbit monoclonal | Connective tissue around portal triad | Abcam/ab215969 |
| Biological samples |  |  |  |  |
| Health mouse liver |  |  |  | 17 weeks old male (n=3) and female (n=3) C57/BL6 mice |
| Health human liver |  |  | Columbia University tissue bank | 51-year-old female |
| Deposited data |  |  |  |  |
| SIMS raw date |  |  | This paper | doi:10.26207/6a38-tr35 |
| DESI raw date |  |  | This paper |  |
| mRNA scope raw date |  |  | This paper |  |
| Software and algorithms |  |  |  |  |
| Ionoptika Image Analyzer |  |  | Ionoptika Ltd. | https://ionoptika.com |
| ImagingSIMS |  |  | Jay Tarolli | https://github.com/ImagingSIMS/ImagingSIMS |
| DeepCell |  |  | Bryan Chan, et al | https://deepcell.com/ |
| SCiLS |  |  | Bruker Corp. | https://www.bruker.com/ |
| Progenesis |  |  | Waters Corp. | https://www.waters.com/ |
| MS<sup>E</sup> dataviewer |  |  | Waters Corp. | https://www.waters.com/ |
| Other |  |  |  |  |

