## Supplemental figures and Tables for "Multi-modal mass spectrometry imaging reveals single-cell metabolic states in mammalian liver"

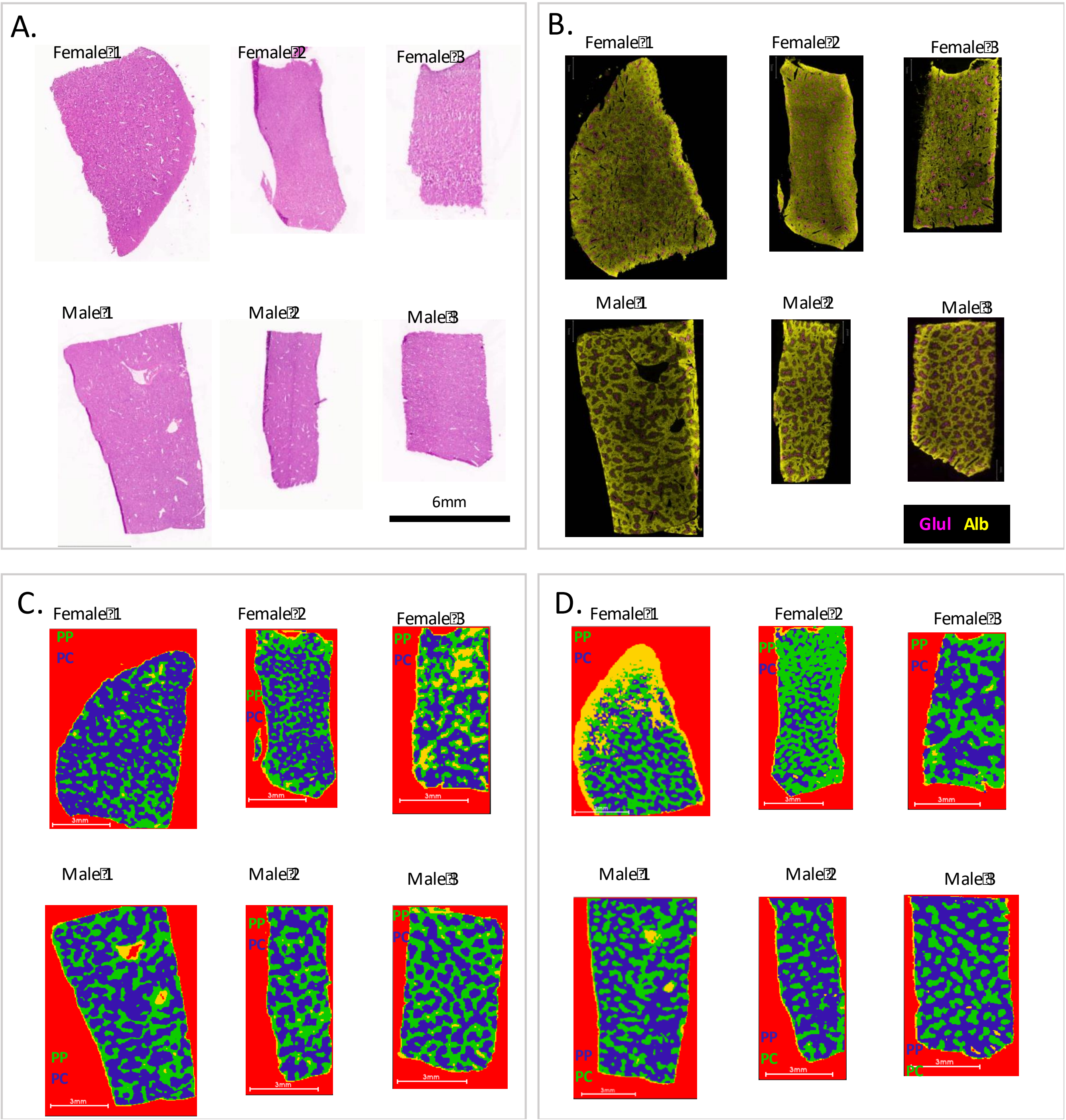


**Figure S1 Identification of pericentral and periportal regions in mouse liver tissues.** (a) Histology of mouse liver sections from 3 female and 3 male mice, assessed by H&E staining (b) RNAScope fluorescence imaging, detecting Glutamine Synthetase (*Glul* -magenta) and Albumin (*Alb* -yellow) RNA transcripts in the corresponding liver tissue sections (c, d) DESI-MSI lipid and metabolite based clustering of pixels within the image, where the clusters corresponding to periportal (PP) and pericentral (PC) regions are denoted by specific colors for data acquired in (c) negative ionization mode and (d) positive ionization mode respectively, in the corresponding liver sections.

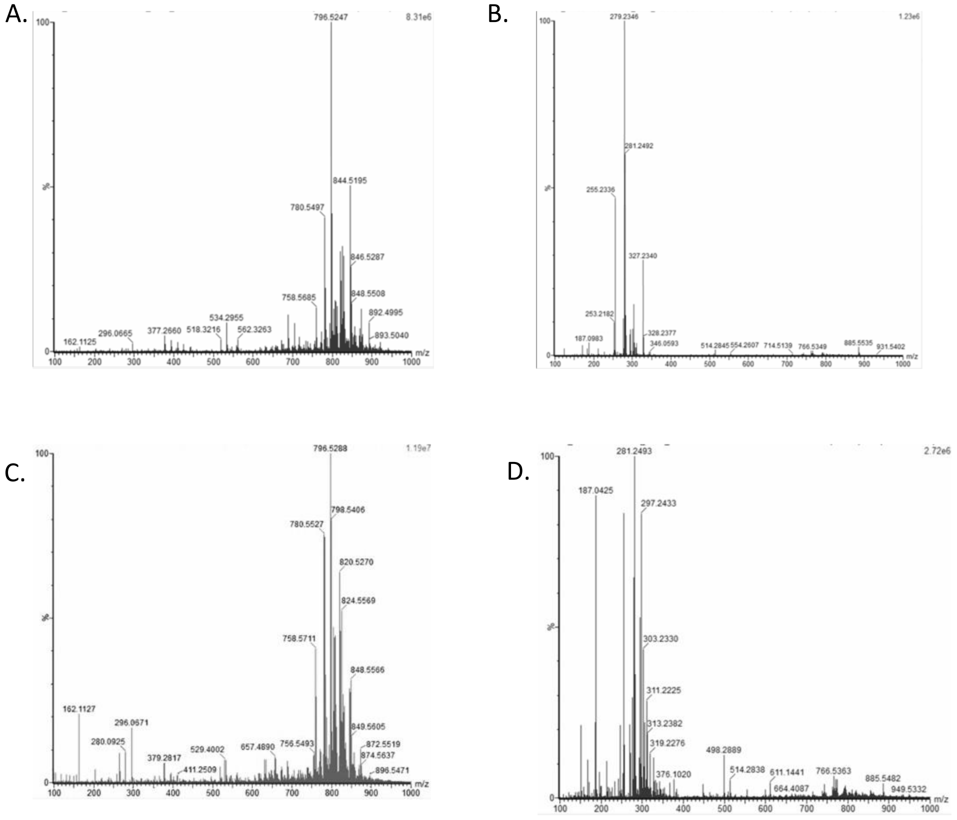


**Figure S2** Representative DESI MSI average mass spectra obtained from (a,b) mouse liver section and (c,d) human liver section, in (a,c) positive and (b,d) negative ion mode.

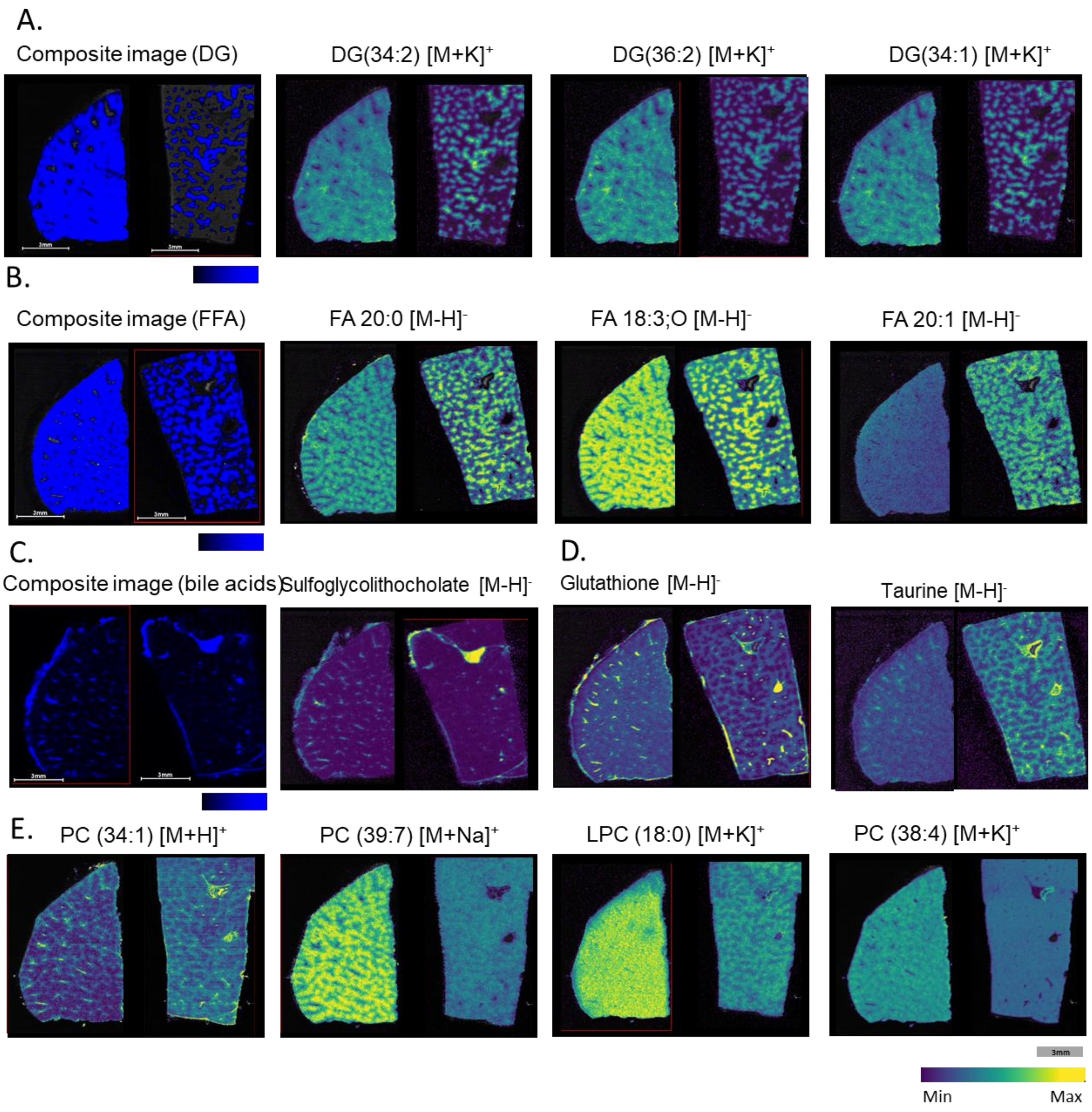


**Figure S3 Representative images of selected metabolites and lipids from DESI MSI of a mouse liver section**. (a) Composite image of diacylglycerols (DG 34:2, DG 34:1, DG 36:2, DG 36:3, DG 36:4, DG 37:7) distribution overlaid with optical image and individual ion images (b) Composite image of free fatty acids (FA 18:1, FA 18:2;O, FA 18:3;O, FA 20:1, FA 16:0, FA 18:2, FA 20:0, FA 20:2, FA 22:5) distribution overlaid with optical image and individual ion images (c) Composite image of conjugated bile acids and an individual ion image (d) Individual images of small metabolites and (e) phosphatidylcholine lipid species by DESI.

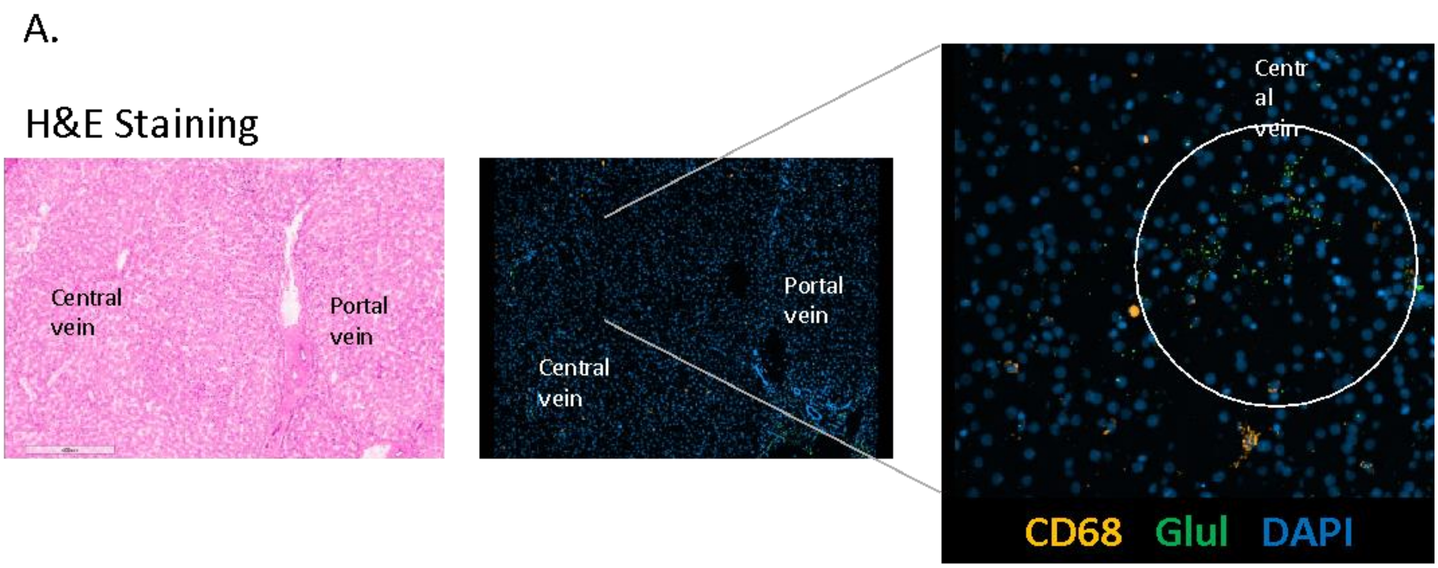


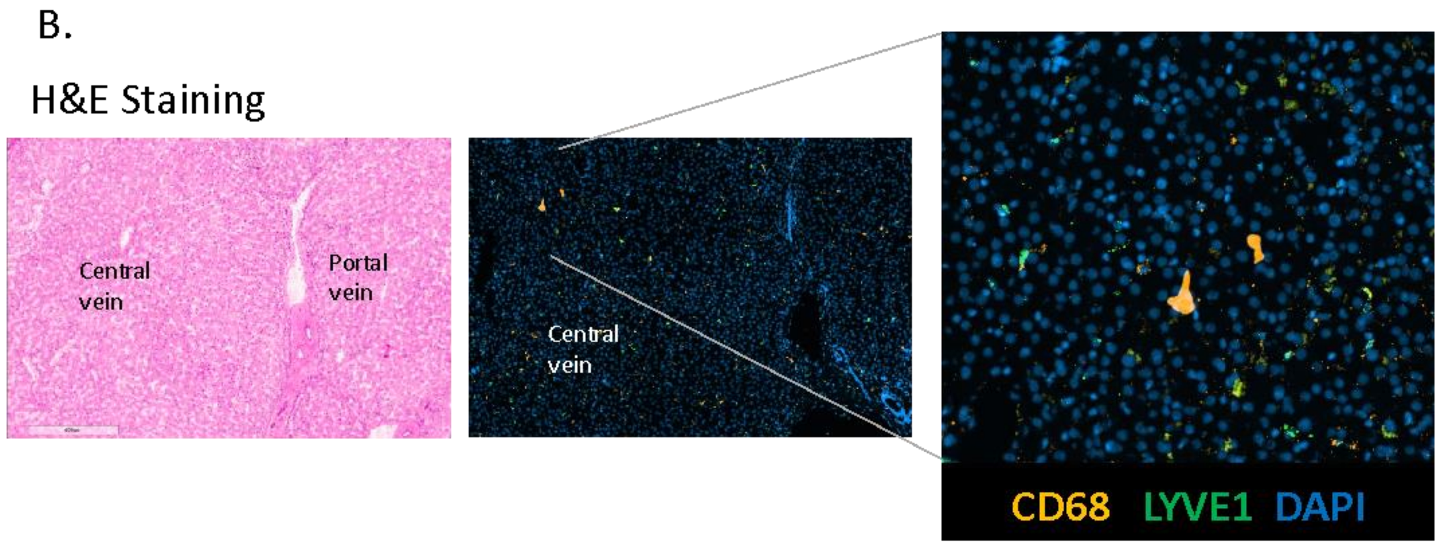


**Figure S4 Fluorescent RNAScope image of cell markers in human liver.**
H&E staining and corresponding RNAScope image on the serial section showing distribution of (a) macrophages (CD68) in yellow, Glutamine Synthetase (Glul) in green and nuclear staining (DAPI) in blue (b) macrophages (CD68) in yellow, sinusoidal endothelial cells (LYVE1) in green and nuclear staining (DAPI) in blue.

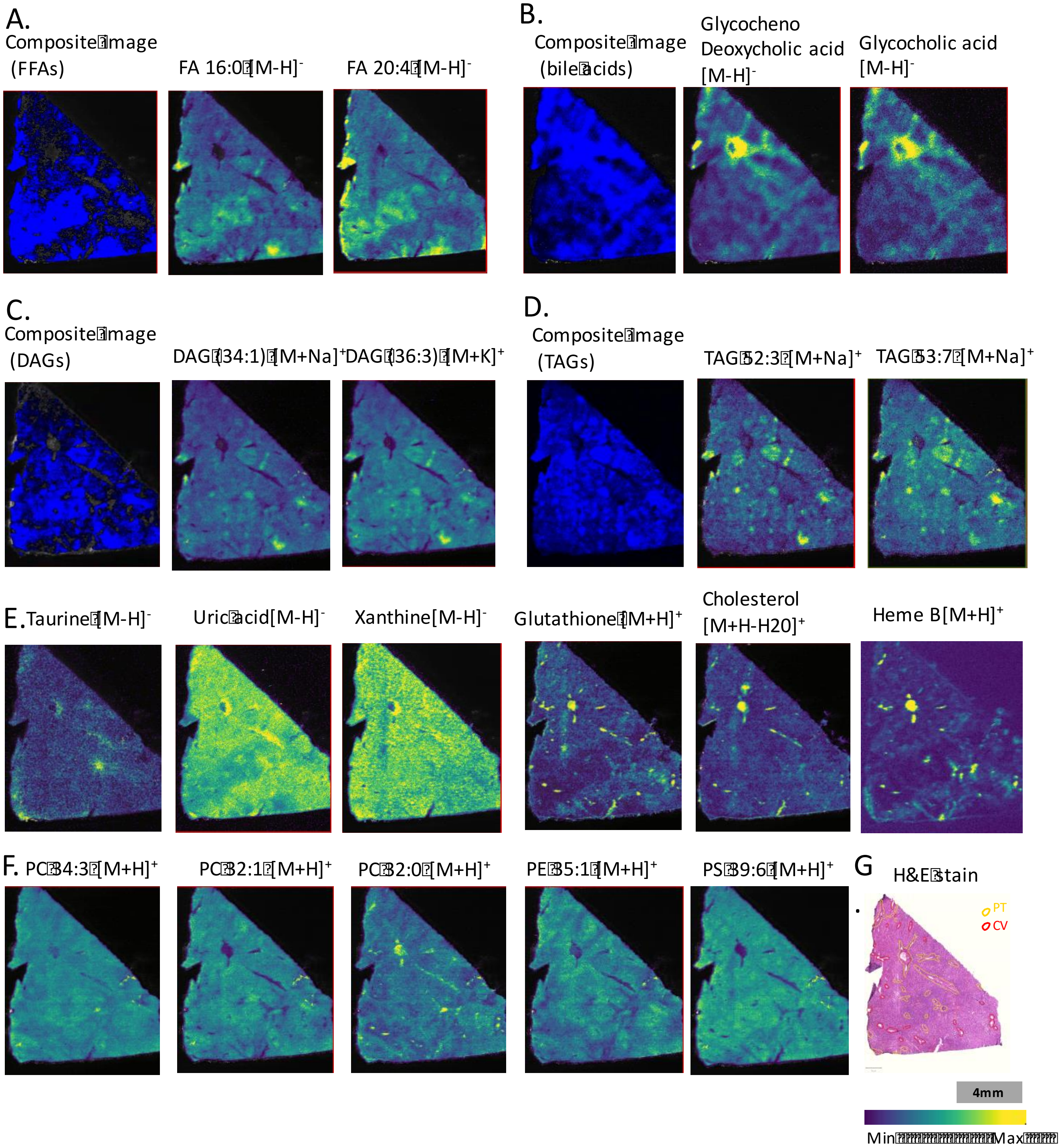


**Figure S5 Representative images of selected metabolites and lipids from DESI MSI of a human liver section** (a) Composite image of free fatty acids FA 18:1, FA 18:2, FA 18:3;O, FA 18:2;O, FA 20:4, FA 20:5 and FA 22:6 overlaid on optical image and individual ion images (b) Composite image of bile acids glycochenodeoxycholic acid, glycocholic acid, taurodeoxycholic acid and taurocholic acid overlaid on optical image, and individual ion images. (c) Composite image of diacylglycerides DAG 34:2, DAG 34:1, DAG 36:4, DAG 36:2, and DAG 36:3, overlaid on optical image, and individual ion images (d) Composite image of triacylglycerides TAG 52:3, TAG 52:2, TAG 53:7, TAG 54:8, TG 55:9, TG 55:8 and TG56:9 overlaid on optical image, and individual ion images (e) Individual ion images of small metabolites (f) Individual ion images of phospholipids (g) H&E staining image for the corresponding DESI images with manual annotation for portal triad (PT) and central vein (CV) region.

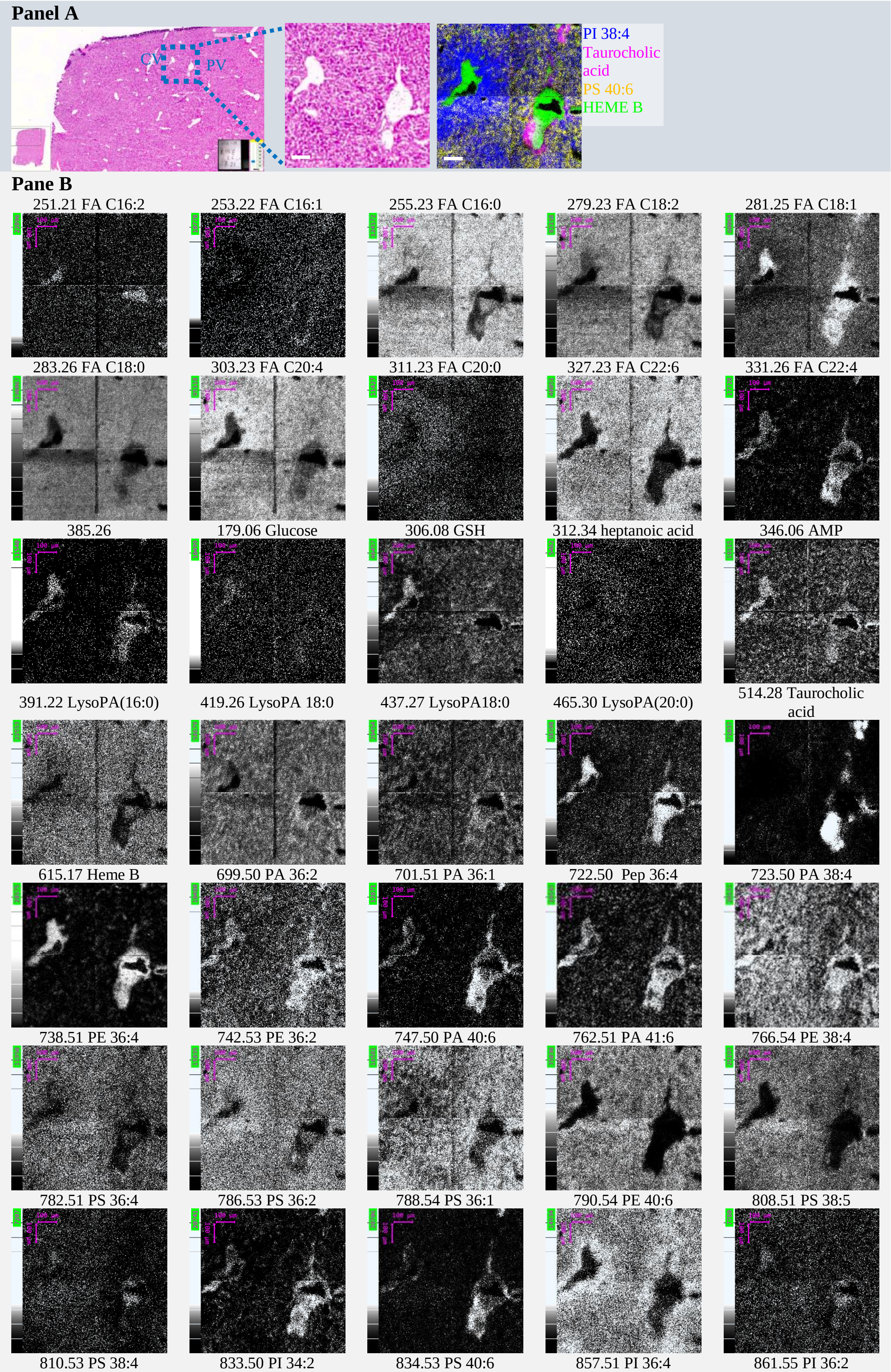
Panel A
CV
PV
PI 38:4
Taurocholic acid
PS 40:6
HEME B
Pane B
251.21 FA C16:2
253.22 FA C16:1
255.23 FA C16:0
279.23 FA C18:2
281.25 FA C18:1
283.26 FA C18:0
303.23 FA C20:4
311.23 FA C20:0
327.23 FA C22:6
331.26 FA C22:4
385.26
179.06 Glucose
306.08 GSH
312.34 heptanoic acid
346.06 AMP
391.22 LysoPA(16:0)
419.26 LysoPA 18:0
437.27 LysoPA18:0
465.30 LysoPA(20:0)
514.28 Taurocholic acid
615.17 Heme B
699.50 PA 36:2
701.51 PA 36:1
722.50 Pep 36:4
723.50 PA 38:4
738.51 PE 36:4
742.53 PE 36:2
747.50 PA 40:6
762.51 PA 41:6
766.54 PE 38:4
782.51 PS 36:4
786.53 PS 36:2
788.54 PS 36:1
790.54 PE 40:6
808.51 PS 38:5
810.53 PS 38:4
833.50 PI 34:2
834.53 PS 40:6
857.51 PI 36:4
861.55 PI 36:2

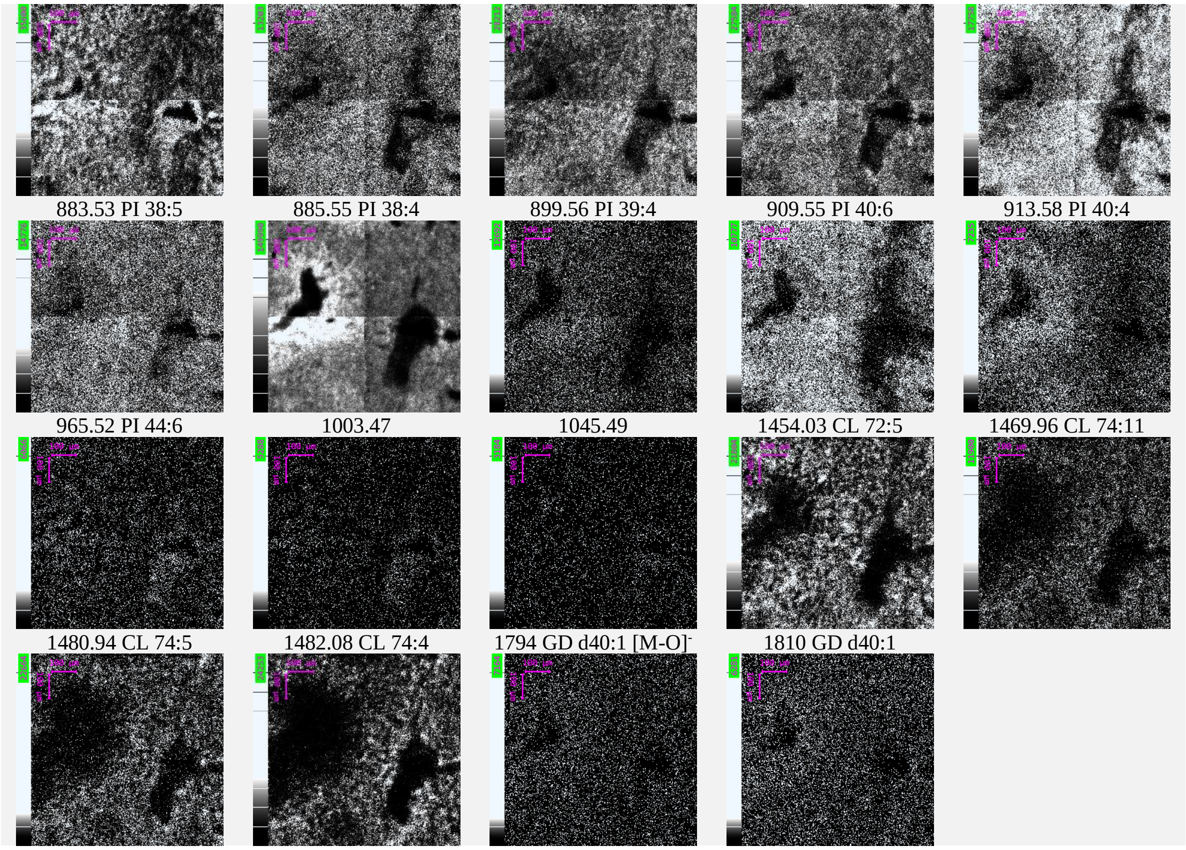


**Figure S6 H&E image and $(H_2O)_n$-GCIB SIMS images of mouse liver section.** Panel A, H&E image of mouse liver tissue on a serial section. An area with the central vein (CV) and portal vein(PV) highlighted in blue, 800×800 $\mu m^2$, was subjected to the $(H_2O)_n$-GCIB SIMS imaging. The color overlay SIMS image shows the anatomic structures by different lipids and proteins. PI 38:4 in Blue is concentrated around the CV, taurocholic acid in magenta is around the PV, PS 40:6 in yellow has elevated intensity around PV, HEME B in blue is mainly inside of the CV and PV. Panel B, selected single ion images of $(H_2O)_n$-GCIB SIMS. If not specified, the species are identified as deprotonated ions $[M-H]^-$. The highly heterogeneous distributions are demonstrated by a variety (>150) of metabolites, lipids and protein, represented by 57 species here. Scale bars 100 µm.

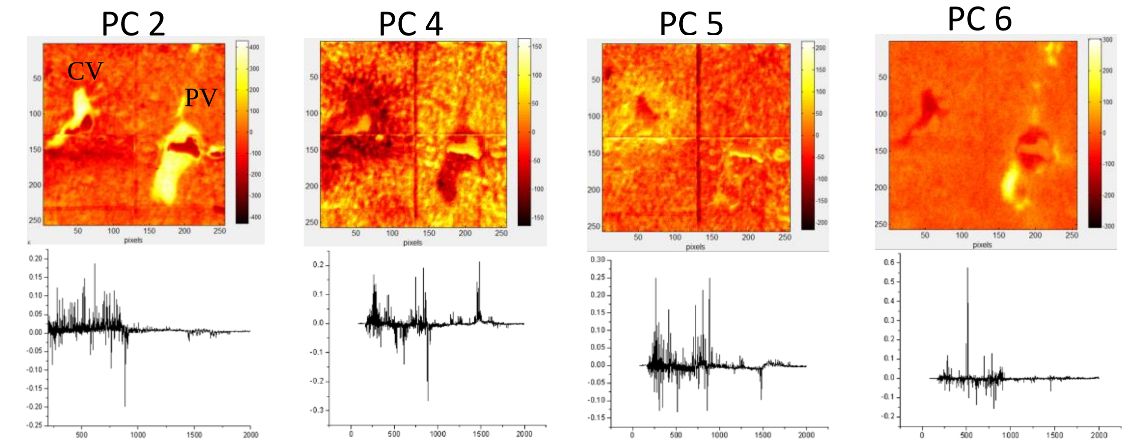


| Top positive loadings | | | | | | | |
|---|---|---|---|---|---|---|---|
| **PC2** | | **PC4** | | **PC5** | | **PC6** | |
| 615.17 | Heme B | 1482.08 | CL 74:5 | 885.55 | PI 38:4 | 514.28 | Taurocholic acid |
| 527.19 | unidentified | 834.53 | PS 40:6 | 266.90 | unidentified | 515.30 | Taurocholic acid (isotopic peak) |
| 512.16 | unidentified | 1482.08 | CL 74:4 | 810.53 | PS 38:4 | 498.30 | Taurodeoxycholic acid/Taurodeoxycholic |
| 616.15 | Heme B (isotopic peak) | 266.90 | unidentified | 268.90 | unidentified | 516.29 | Taurocholic acid |
| 281.25 | FA C18:1 | 747.50 | PA 40:6 | 886.59 | unidentified | 512.26 | Sulfoglycolithocholate |
| 786.52 | PS 36:2 | 1454.92 | unidentified | 723.50 | PA 38:4 | 788.54 | PS 36:1 |
| 701.51 | PA 36:1 | 835.50 | PI 34:1 | 419.26 | LysoPA 18:0 | 283.26 | FA C18:0 |
| 512.26 | Sulfoglycolithocholate | 283.26 | FA C18:0 | 811.53 | PS 38:4 | 701.51 | PA 36:1 |
| 750.54 | PEp(38:4) | 268.90 | unidentified | 303.23 | FA C20:4 | 512.16 | unidentified |
| 514.28 | Taurocholic acid | 1484.98 | unidentified | 812.54 | PS 38:3 | 789.55 | PS 36:1 |
| | | 1455.95 | CL 72:4 | | | | |
| | | 1468.95 | unidentified | | | | |
| | | 1480.9 | unidentified | | | | |

**Figure S7 PCA analysis of mouse liver tissue image using $(H_2O)_n$-GCIB SIMS.** The scores and loadings of PC 2-6 show the lipid/metabolites species that distinguish the major features around the CV and PV. The top species contributing to the positive loading are listed in the tables.

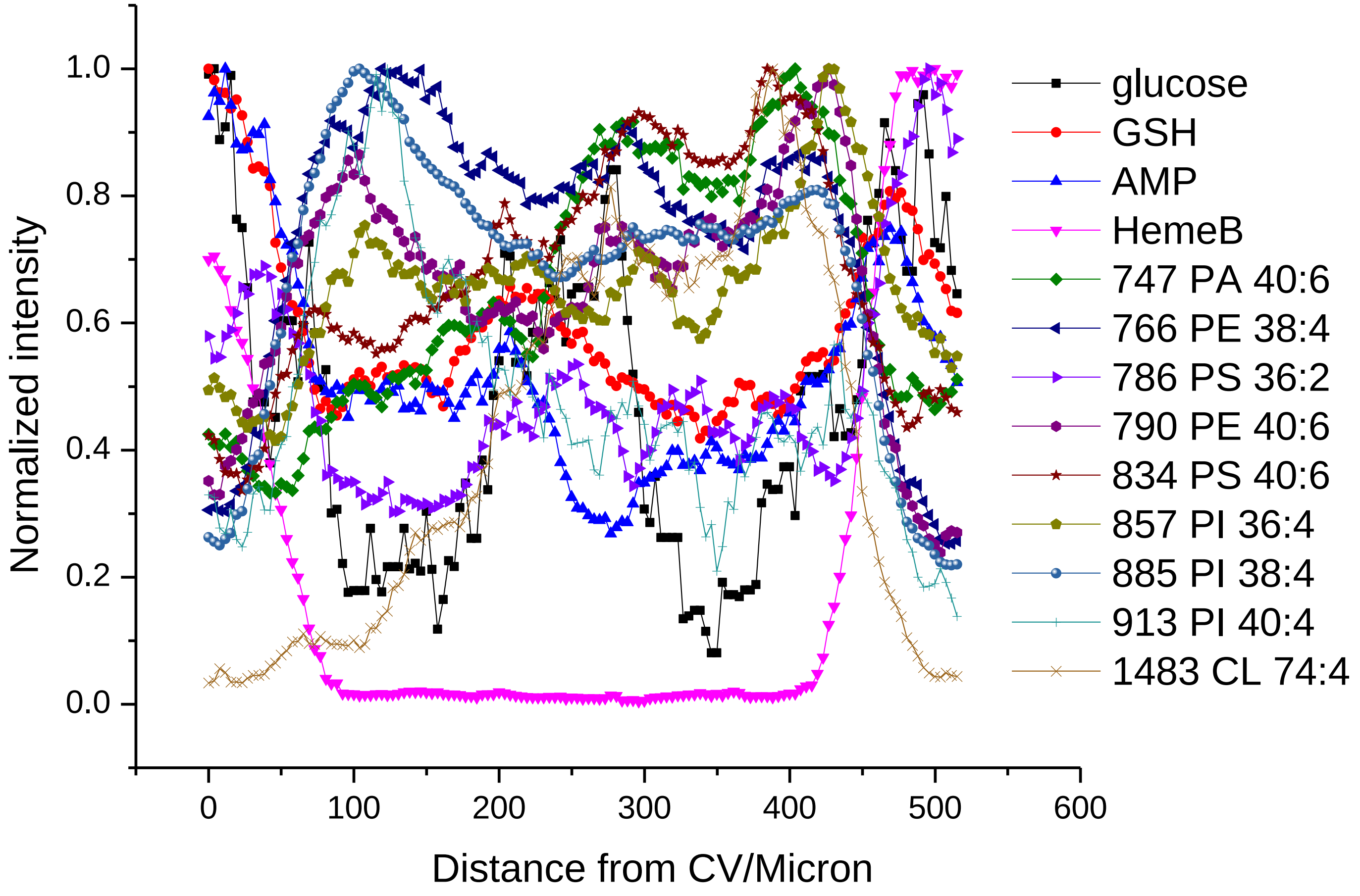


**Figure S8** Variation of various species from CV to PV on mouse liver tissue.

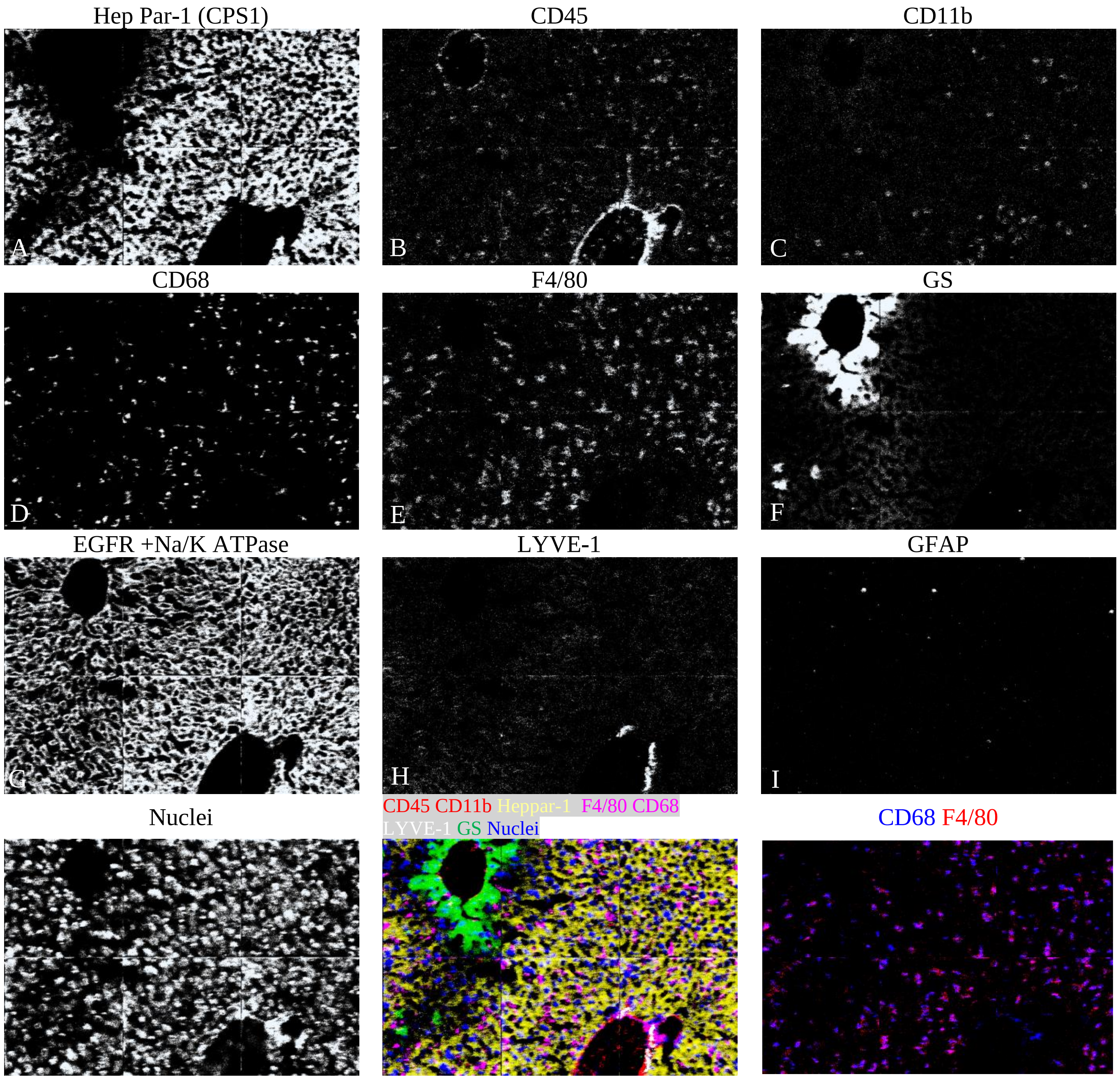


**Figure S9** Multiplexed imaging of lanthanide-tagged antibodies on CV and PV region of mouse liver tissue using $C_{60}$-SIMS. The antibodies panel is detailed in Table S1. The single channel images of each antibody are shown is A-J. Color overlay image K shows the distribution of CD 45 and CD11b in red, Heppar-1 in yellow, F4/80 and CD68 in magenta, LYVE-1 in white, GS in Green and nuclei in blue. Color overlay image L shows the distribution of CD 68 in blue and F4/80 in red, with the overlapping area in magenta.

**Panel A**

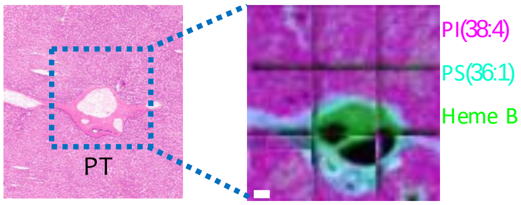


**Panel B**

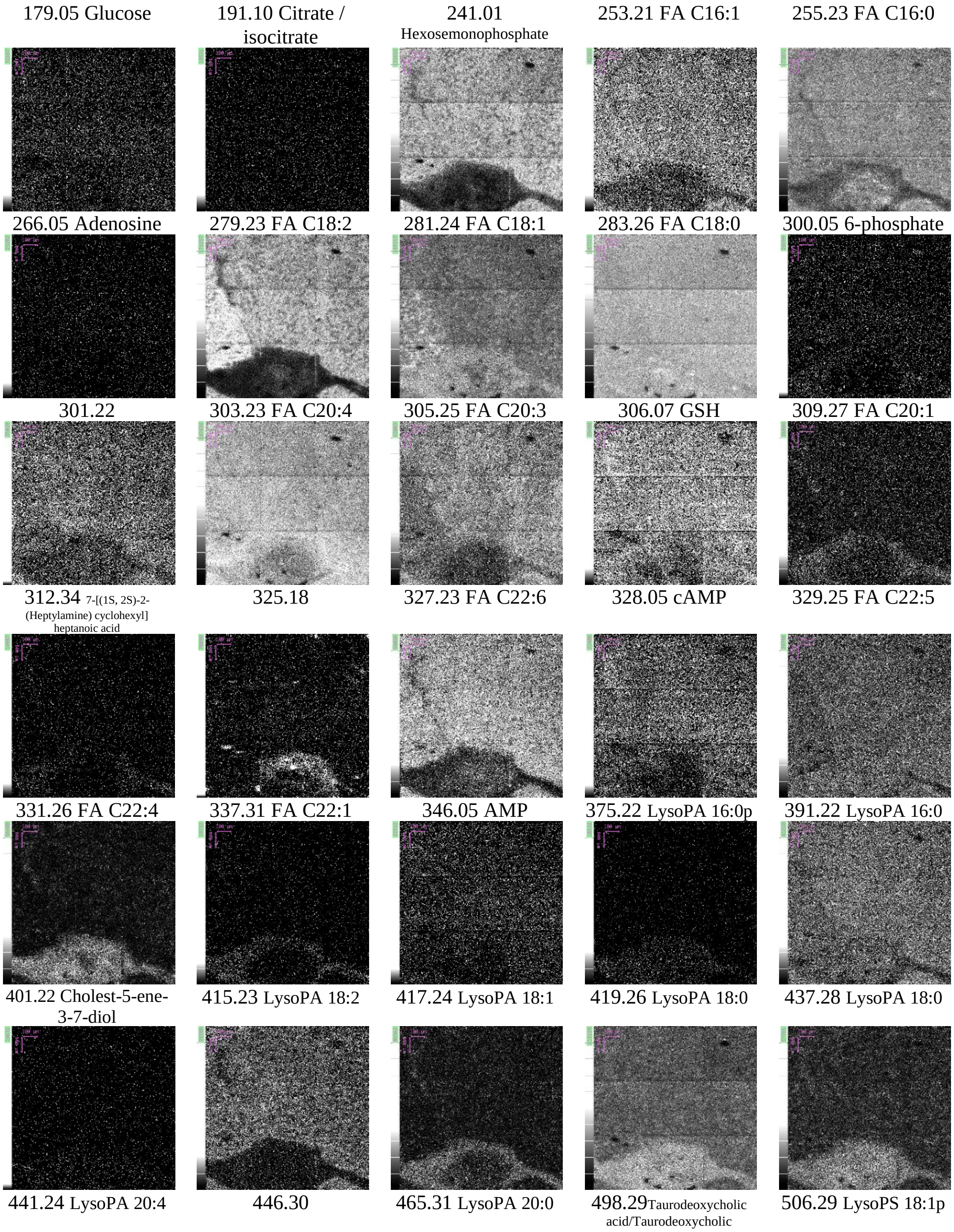

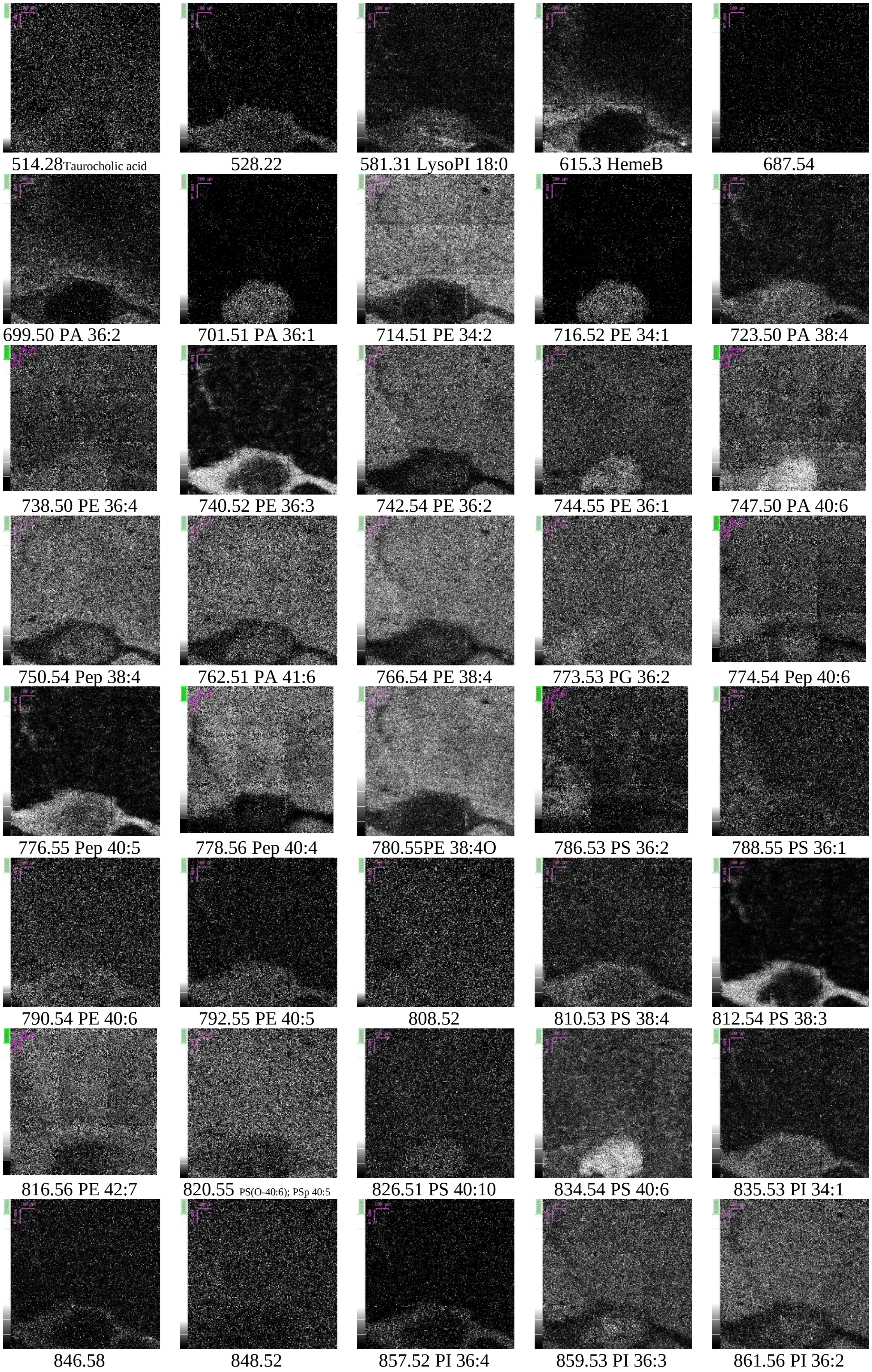

514.28 Taurocholic acid
528.22
581.31 LysoPI 18:0
615.3 HemeB
687.54
699.50 PA 36:2
701.51 PA 36:1
714.51 PE 34:2
716.52 PE 34:1
723.50 PA 38:4
738.50 PE 36:4
740.52 PE 36:3
742.54 PE 36:2
744.55 PE 36:1
747.50 PA 40:6
750.54 Pep 38:4
762.51 PA 41:6
766.54 PE 38:4
773.53 PG 36:2
774.54 Pep 40:6
776.55 Pep 40:5
778.56 Pep 40:4
780.55PE 38:4O
786.53 PS 36:2
788.55 PS 36:1
790.54 PE 40:6
792.55 PE 40:5
808.52
810.53 PS 38:4
812.54 PS 38:3
816.56 PE 42:7
820.55 PS(O-40:6); PSp 40:5
826.51 PS 40:10
834.54 PS 40:6
835.53 PI 34:1
846.58
848.52
857.52 PI 36:4
859.53 PI 36:3
861.56 PI 36:2

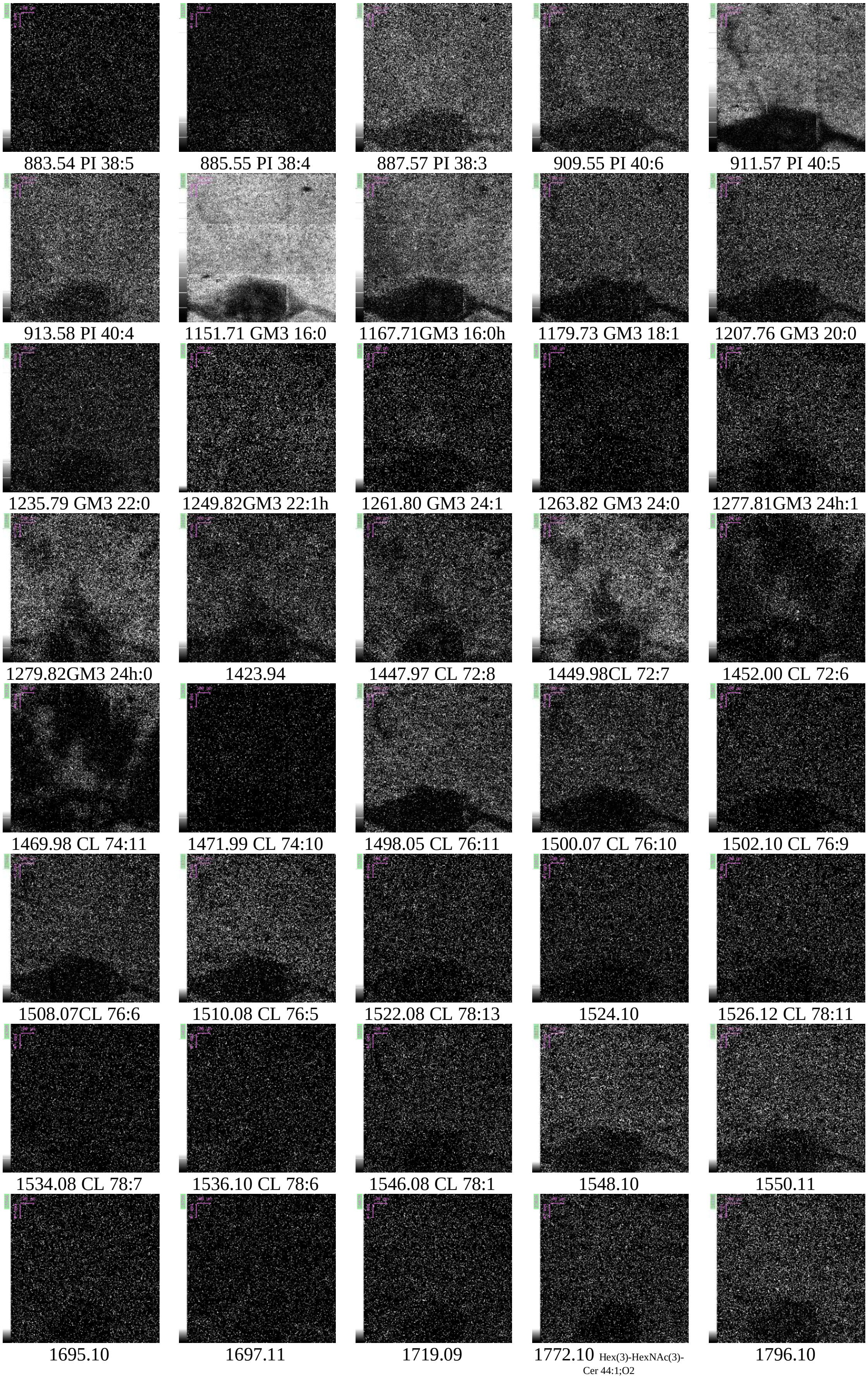
883.54 PI 38:5
885.55 PI 38:4
887.57 PI 38:3
909.55 PI 40:6
911.57 PI 40:5
913.58 PI 40:4
1151.71 GM3 16:0
1167.71GM3 16:0h
1179.73 GM3 18:1
1207.76 GM3 20:0
1235.79 GM3 22:0
1249.82GM3 22:1h
1261.80 GM3 24:1
1263.82 GM3 24:0
1277.81GM3 24h:1
1279.82GM3 24h:0
1423.94
1447.97 CL 72:8
1449.98CL 72:7
1452.00 CL 72:6
1469.98 CL 74:11
1471.99 CL 74:10
1498.05 CL 76:11
1500.07 CL 76:10
1502.10 CL 76:9
1508.07CL 76:6
1510.08 CL 76:5
1522.08 CL 78:13
1524.10
1526.12 CL 78:11
1534.08 CL 78:7
1536.10 CL 78:6
1546.08 CL 78:1
1548.10
1550.11
1695.10
1697.11
1719.09
1772.10 Hex(3)-HexNAc(3)-Cer 44:1;O2
1796.10

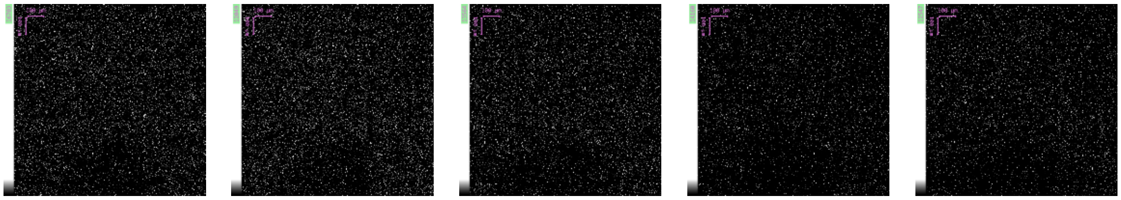

**Figure S10** H&E image and $(H_2O)_n$-GCIB SIMS images of human liver section. Panel A, H&E image of human liver tissue on a serial section. An area with portal Triad (PT) highlighted in blue, 1200×1200 $\mu m^2$, was subjected to the $(H_2O)_n$-GCIB SIMS imaging. The color overlay SIMS image shows the anatomic structures by different lipids and protein, PI 38:4 in magenta is outside of the PT, PS 36:1 in blue is around the PT, HEME B in blue is mainly inside of the PT. Panel B, selected ion images of $(H_2O)_n$-GCIB SIMS. The highly heterogeneous distribution are demonstrated by a variety (>150) of metabolites, lipids and protein.

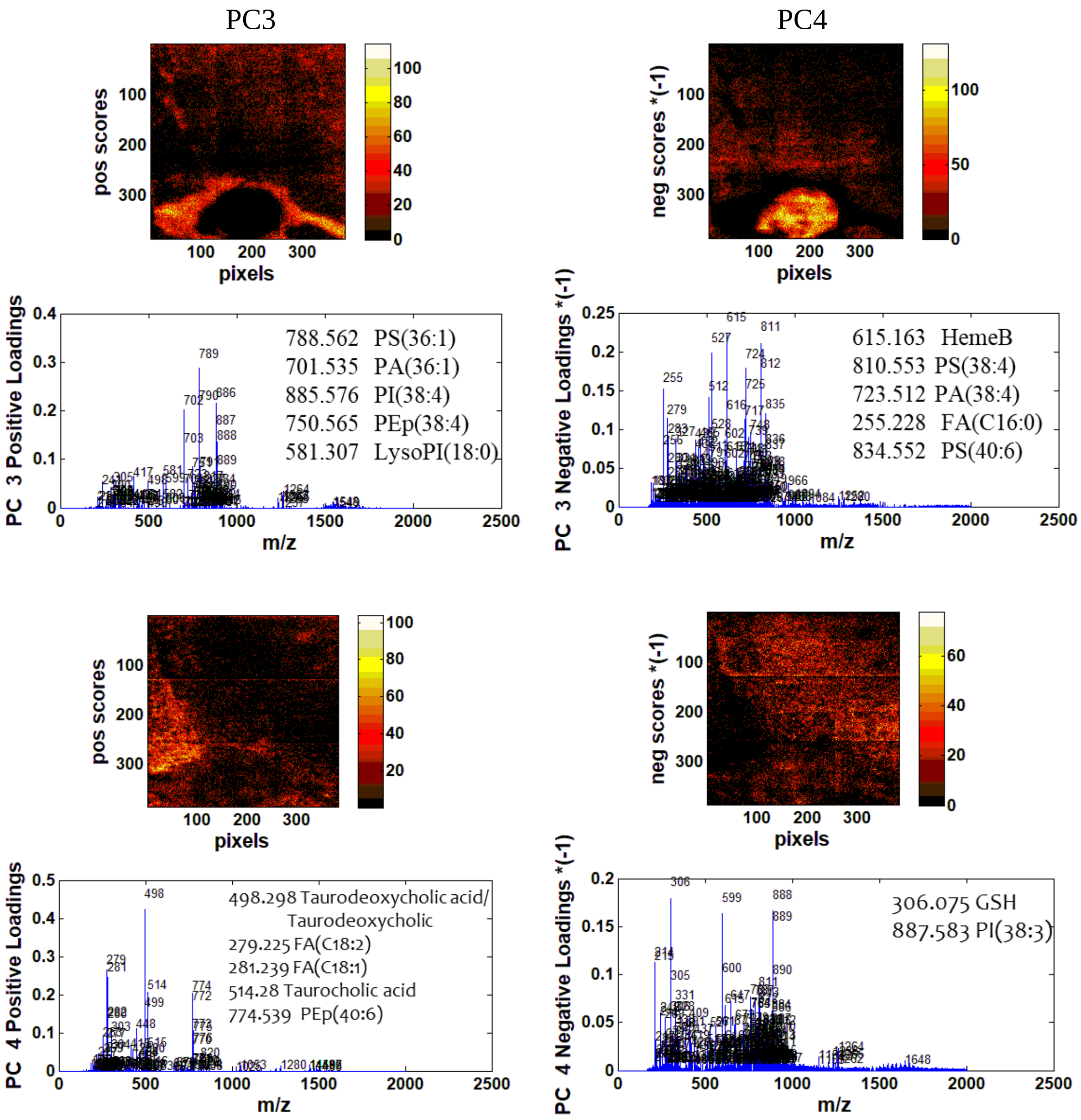


**Figure S11** PCA analysis of human liver tissue image using $(H_2O)_n$-GCIB SIMS. The positive and negative scores and loadings of PC 3 and 4 show the lipid/metabolites species that distinguish the major features and metabolite flux around the PT.

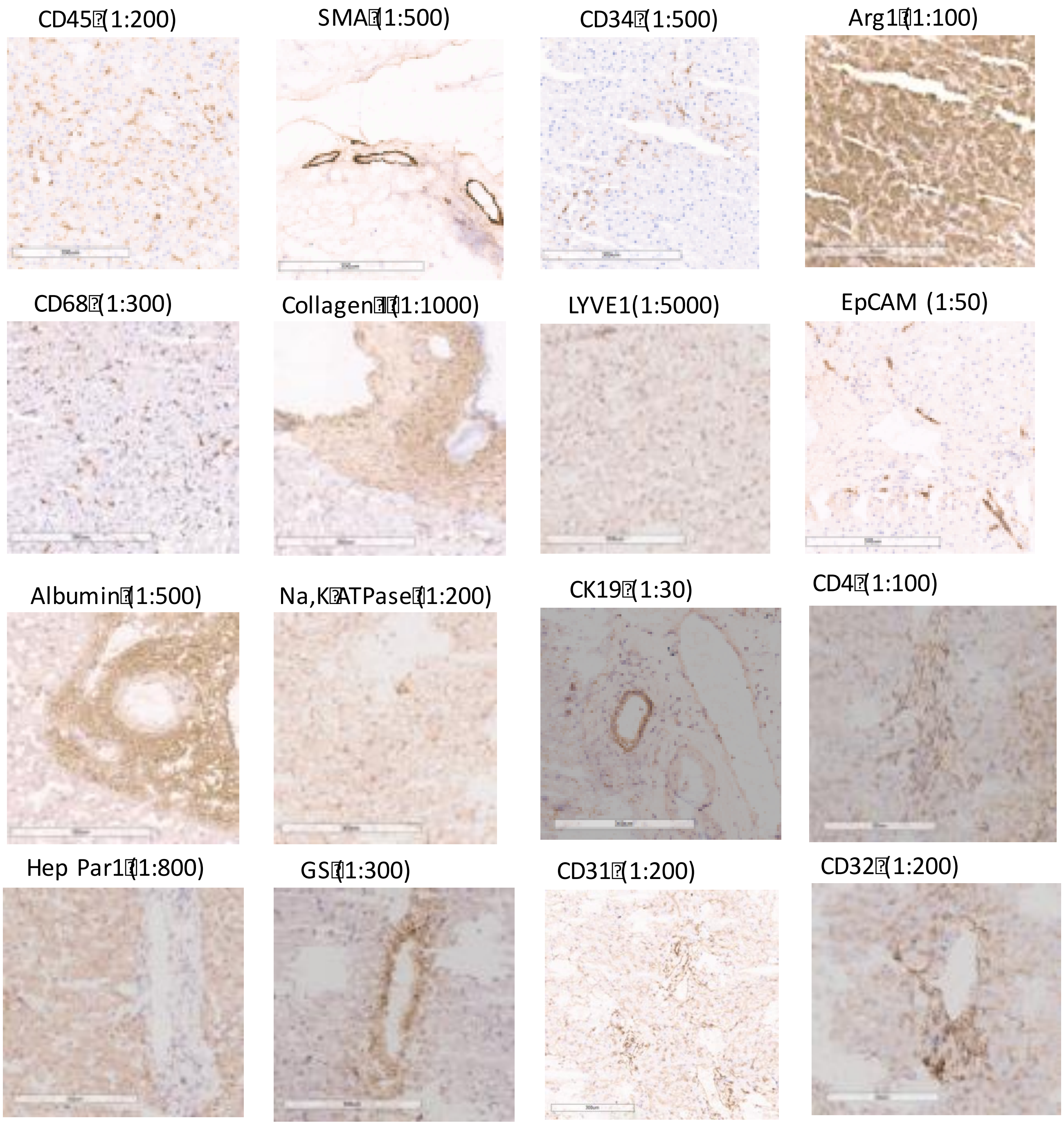


**Figure S12**. The panel of antibodies were validated for specificity and sensitivity by immunohistochemistry in human liver tissue section.

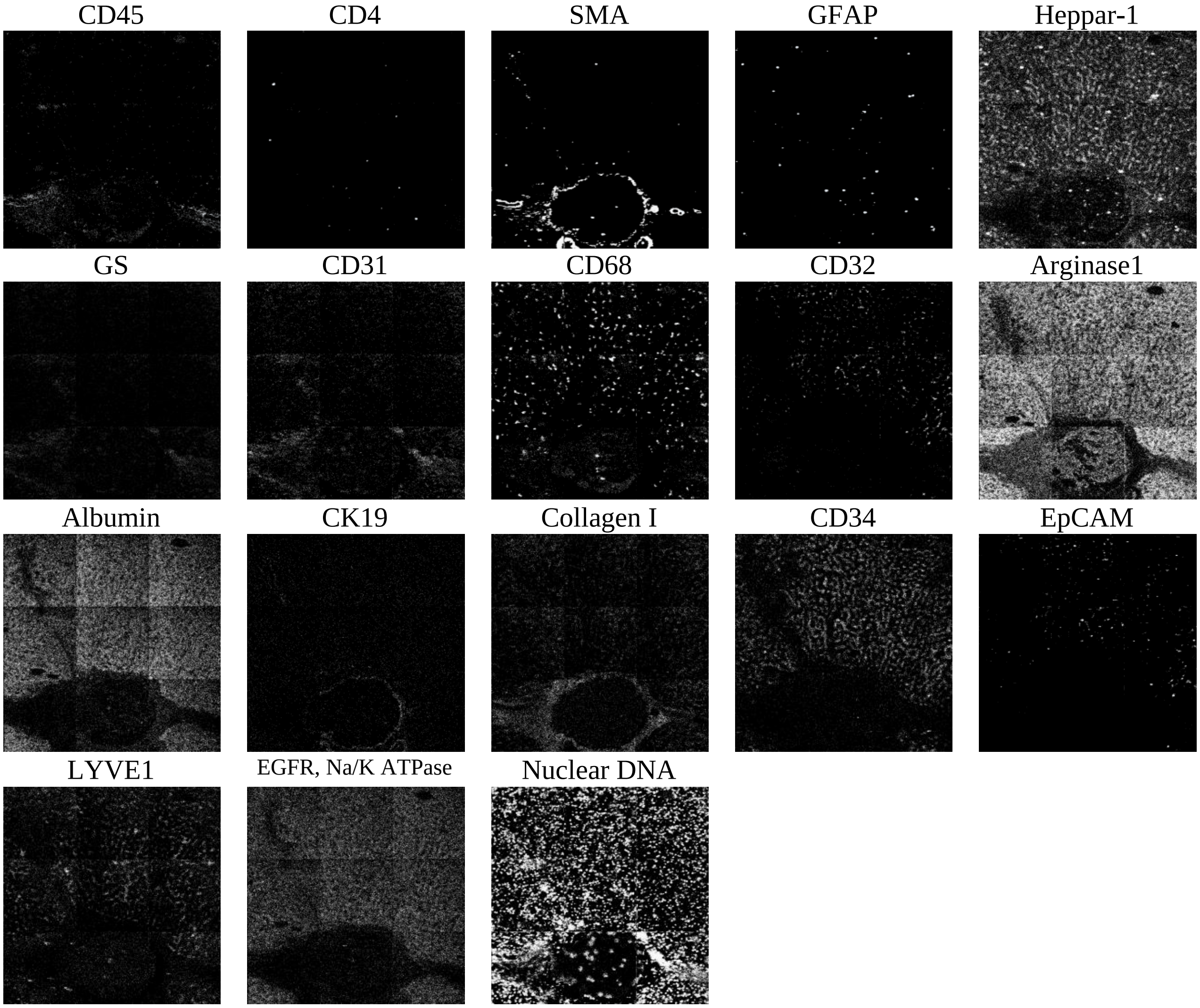


**Figure S13** Multiplexing imaging of lanthanides-tagged antibodies on portal triad region of human liver tissue using $C_{60}$-SIMS. The antibodies panel is detailed in Table S2. Because the central vein is out of the field of view of $C_{60}$-SIMS image, GS is not detected.

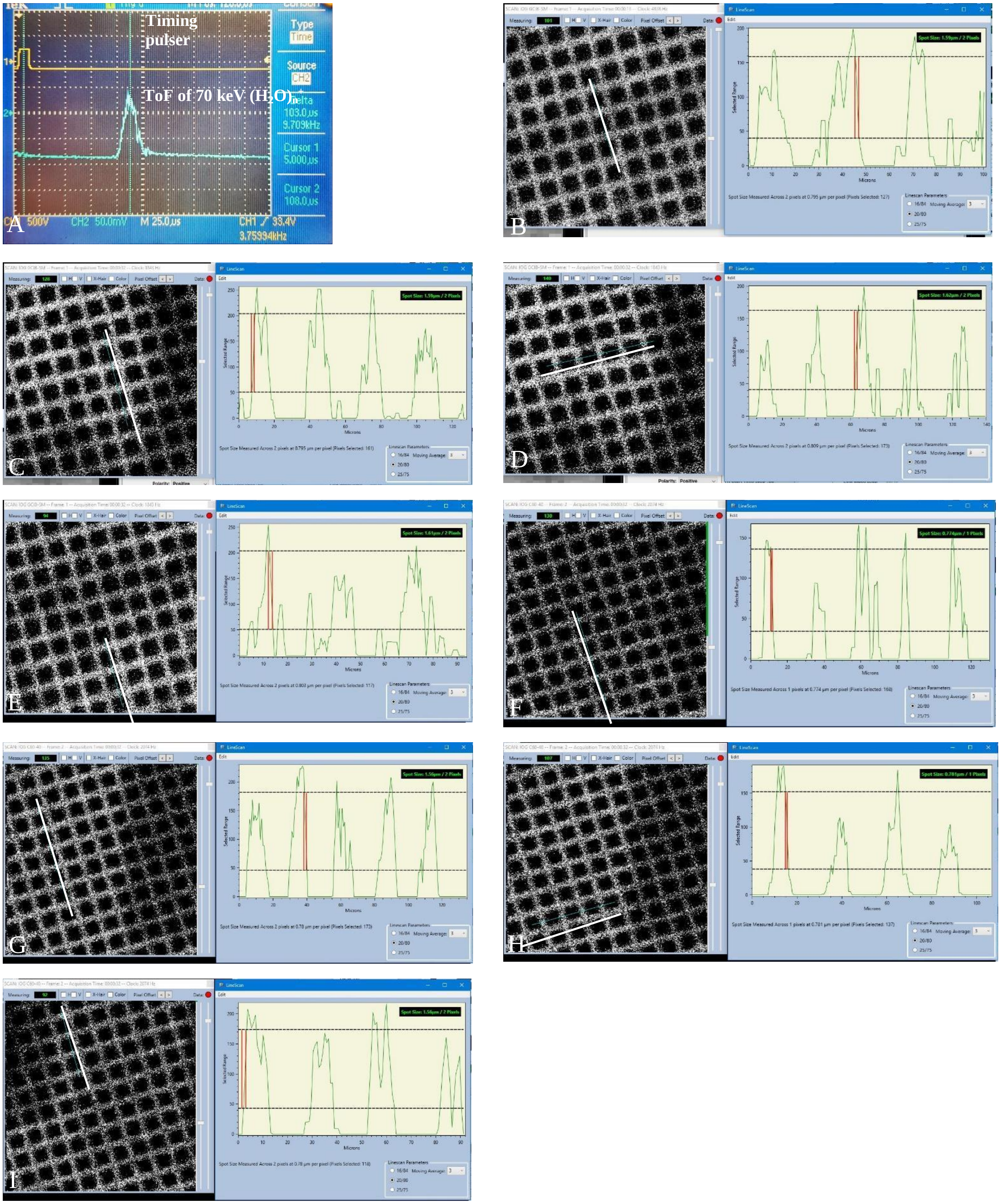


**Figure S14 Time-of-Flight (ToF) and beam spot size measurements**. of 70 keV $(H_2O)_{30k}^+$ and 40 kV $C_{60}^+$. A: Oscilloscope reading of the timing pulse (upper line) and $(H_2O)_n$ cluster beam (lower line). The measured ToF of the cluster is 103 µs with a peak width at half maximum of 13 µs. The beam energy is 70 kV, distance to the sample surface is 0.533 m, and the SED offset is 8 µs. Using these values gives a calculated n = 30,900 for the 70 keV $(H_2O)_{30k}^+$ GCIB. B-E, 70 keV $(H_2O)_{30k}^+$ focus measurements by line scanning across the bars of an SED image of a 1000 mesh grid. The average beam spot is 1.60±0.01 µm (mean $\pm$ std dev of the measurements in B-E). F-I, 40 kV $C_{60}^+$ focus measurements by line scanning across the bar of SED image of grid (1000 mesh). The average beam spot is 1.16±0.45 µm (mean $\pm$ std dev of the measurements in F-I)

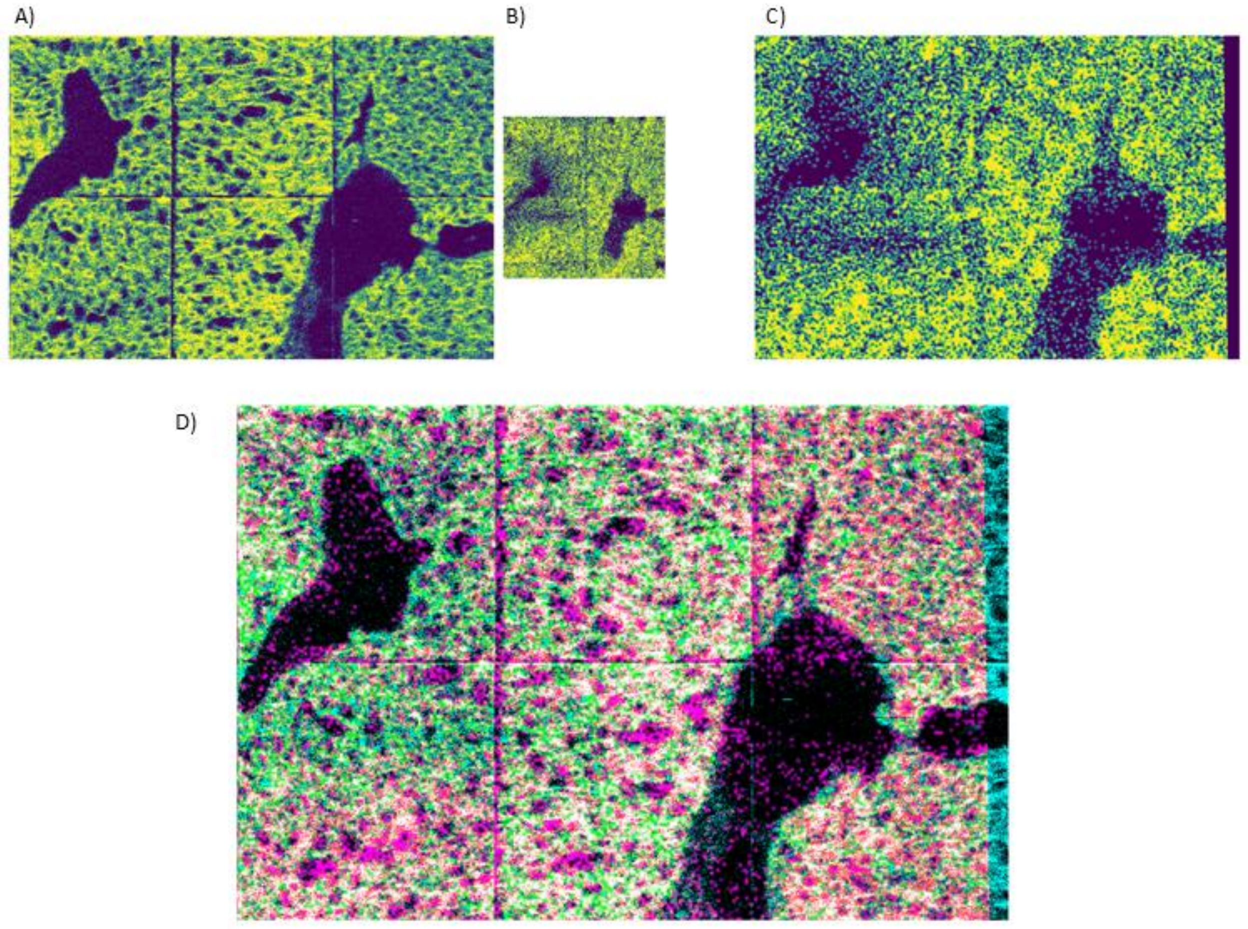


**Figure S15** Co-registration of $C_{60}$ and $(H_2O)_n$-GCIB SIMS images of mouse liver: A) EGFR ATPase $C_{60}$ -SIMS image (512 x 768 pixels) B) sum of *m/z* 1455 (CL 72:5), *m/z* 1643 (not assigned), and *m/z* 1483 (CL 74:4) $(H_2O)_n$-GCIB SIMS images (256 x 256 pixels, C) $(H_2O)_n$-GCIB SIMS image from (B) with the registration transform applied, rescaled into the image space of (A), D) overlay of C (magenta) and A (cyan) showing the registration of the $(H_2O)_n$-GCIB SIMS image to the image space of the C60 image. Co-registration of $C_{60}$ and $(H_2O)_n$-GCIB SIMS images was performed in Python using SimpleITK (v 2.0.2). First, m/z windows for each SIMS image set were chosen that well represented the morphology of the tissue and were comparable in signal distribution. Because $C_{60}$ -SIMS total ion images in could contain signal from tissue-free regions of the sample, the morphology may appear different in the $C_{60}$ -SIMS and $(H_2O)_n$-GCIB SIMS images, in which case an individual mass image of a generic membrane tag was selected for registration. For $(H_2O)_n$-GCIB SIMS images, the sum of two or three lipid images was used for registration, defined as the moving image. Both registration images were subjected to thresholding to remove low count background signal and normalization. The moving image was registered to the image space of the fixed image ($C_{60}$ -SIMS total ion or membrane tag) using gradient descent optimization of the mean square difference between each image. An initial affine transform was estimated to roughly align the two images before optimization. The result of co-registration of the mouse liver sample is shown in D). The optimized affine transform was then applied to all extracted m/z windows from the $(H_2O)_n$-GCIB SIMS data set.

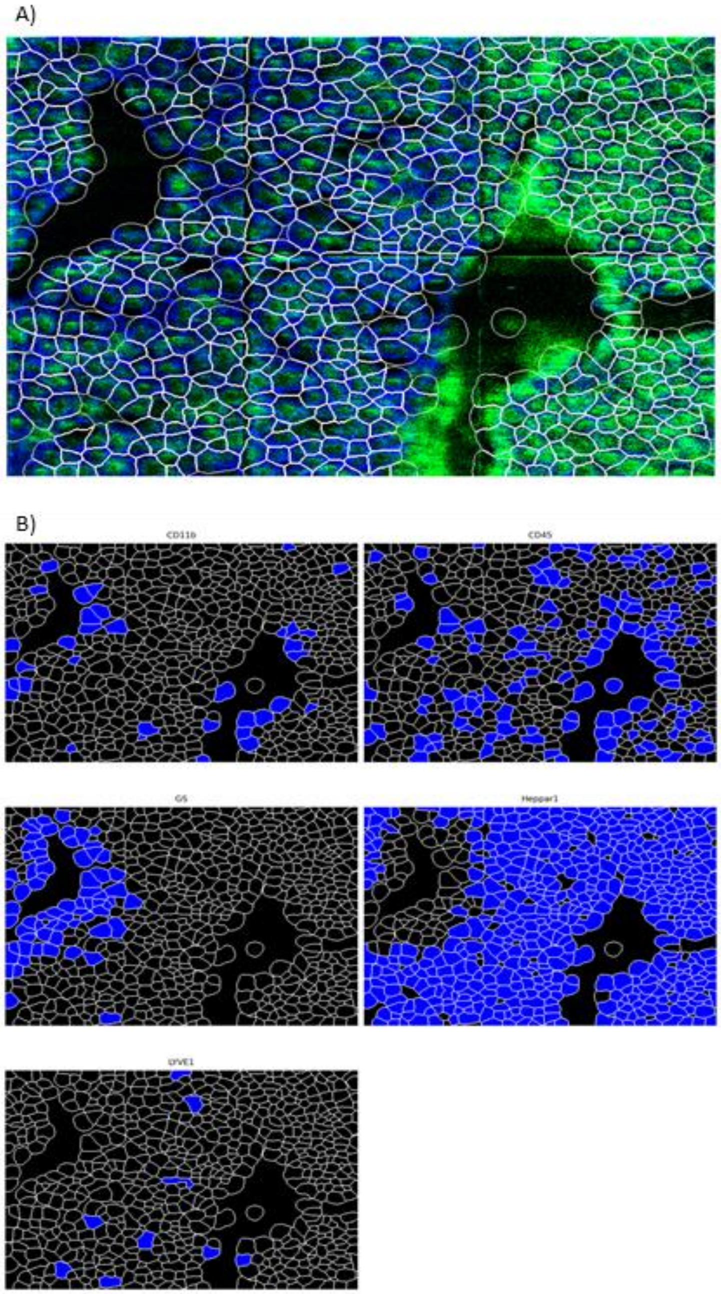


**Figure S16** A) Cell boundaries (white) determined by Mesmer overlayed with the nuclear (green) and membrane (blue) tags, B) Classified cell types for each of the other protein tags in the $C_{60}$-SIMS data set. The $(H_2O)_n$-GCIB-SIMS images were already transformed into the $C_{60}$ image space, so the cell boundaries determined by Mesmer will be applicable to both data sets. The signal intensity in each cell instance was integrated for each lipid and metabolite species of interest. Hierarchical clustering analysis (HCA) was performed with seaborn (v 0.11.1) on the complete data set of cell types determined from the $C_{60}$ images and the integrated lipid and metabolite signal from the $(H_2O)_n$-GCIB images. Whole cell segmentation was performed with Mesmer, an algorithm included in DeepCell (v 0.9.0) which segments tissue images into single cell instances using a trained machine learning model. Since the $C_{60}$ images have an inherently better spatial resolution, segmentation was performed using the nuclear and membrane tags from the $C_{60}$-SIMS data set. Cell instances were classified as positive or negative for each of the protein markers from the $C_{60}$-SIMS data by gating on manually determined thresholds.

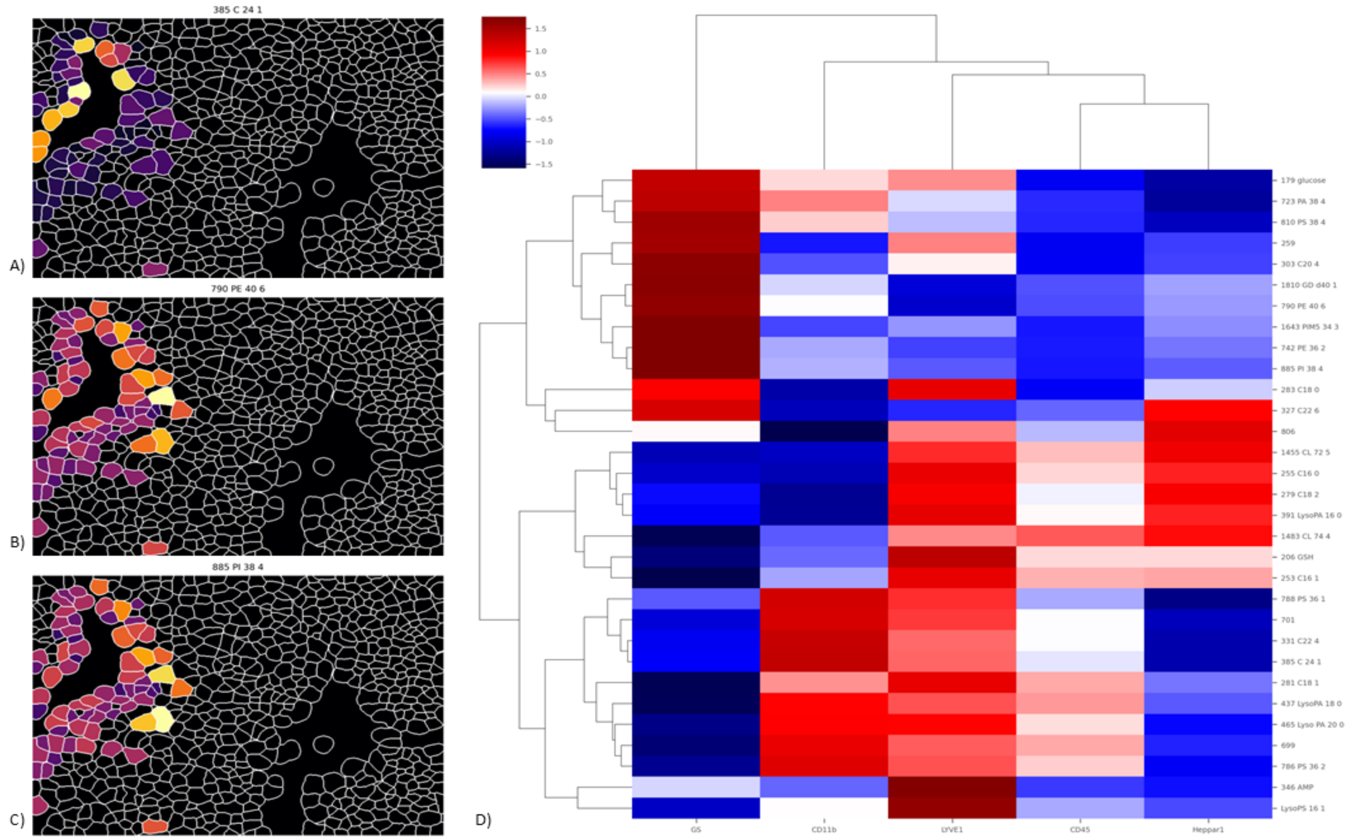


**Figure S17** Expression of three lipids A) *m/z* 385 C 24 1, B) *m/z* 885 PI 38:4, and C) *m/z* 790 PE 40:6 on $GS^+$ cells and D) HCA results plotting the average expression of each lipid or metabolite within cell types.

### Table S1 Antibody panel for mouse liver tissue

| Metal tag | Target | Clone/Host | Cell/Pathway | Manufacturer/Catalog No. |
|---|---|---|---|---|
| **89Y** | CD45 | 30-F11, Rat IgG2b | Pan leukocyte | Fluidigm/3089005B |
| **142Nd** | Heppar-1/CPS1 | HepPar1, mouse monoclonal | Periportal hepatocytes (Zone 1) | Novus/NBP3-08970 |
| **143Nd** | GFAP | EPR1034Y, Rabbit monoclonal | Ito Stellate Cells | Abcam/ab218309 |
| **147Sm** | GS | ab240193,Rabbit monoclonal | Pericentral hepatocytes (Zone 3) | Abcam/ab240193 |
| **152Sm** | F4/80 | CI:A3-1, Mouse IgG2b | Kupffer cells | Abcam/ab6640 |
| **155Gd** | CD11b | M1/70, Rat IgG | Leukocytes | Ionpath/ 715503-100 |
| **156Gd** | CD68 | FA-11, Mouse IgG2b | Macrophages | Abcam/ ab237968 |
| **174Yb** | LYVE1 | Rabbit polyclonal | Sinusoidal endothelial cells | Novus/ NB600-1008 |
| **176Yb** | EGFR | EP38Y, Rabbit monoclonal | Cell membrane | Abcam/ ab272293 |
| **176Yb** | Na/K ATPase | EP1845Y, Rabbit monoclonal | Cell membrane | Abcam/ab167390 |
| **191 Ir** | Nuclear DNA | | Nuclei | Fluidigm/ 201192B |

### Table S2 Antibody panel for human liver tissue

| Metal tag | Target | Clone/Host | Cell/Pathway | Manufacturer/Catalog No. |
|---|---|---|---|---|
| **89Y** | CD45 | D9M8I, Rabbit IgG | Pan leukocyte | Cell Signaling/13917S |
| **113 In** | CD4 | RPA-T4, mouse monoclonal | T cells and macrophages | Novus/NBP2-25199 |
| **141Pr** | SMA | 1A4, Mouse IgG2a | Smooth muscle cells in vascular walls, sinusoildal endothelial cells and activated fibroblasts | Fluidigm/3141017D |
| **143Nd** | GFAP | EPR1034Y, Rabbit monoclonal | Ito Stellate Cells | Abcam/ab218309 |
| **145Nd** | Heppar-1/CPS1 | HepPar1, mouse monoclonal | Periportal hepatocytes (Zone 1) | Novus/NBP3-08970 |
| **147Sm** | GS | ab240193,Rabbit monoclonal | Pericentral hepatocytes (Zone 3) | Abcam/ab240193 |
| **148Nd** | CD31 | JC/70A, Mouse monoclonal | Endothelial cells | Abcam/ab264090 |
| **151Eu** | CD68 | D4B9C,Rabbit IgG | Macrophages | Cell Signaling/76437S |
| **153Eu** | CD32 | FUN-2, Mouse IgG2b | B cells, monocytes, granulocytes and platelets | Fluidigm/3153018B |
| **158Gd** | Arginase1 | D4E3M™,Rabbit IgG | Zone 1-2 hepatocytes, and macrophages | Cell Signaling/93668S |
| **161Dy** | Albumin | EPR20195, Rabbit monoclonal | Periportal hepatocytes (Zone 1) | Abcam/Ab271979 |
| **166Er** | CK19 | SPM561, Mouse monoclonal | Cholangiocytes (Portal triad) | Abcam/ab212569 |
| **169Tm** | CD34 | QBEND/10, Mouse / IgG1 | Endothelial cells | Abcam/ ab198395 |
| **170 Er** | EpCAM | E6V8Y, Rabbit IgG | Hepatic stem/progenitor cells | Cell Signaling /93790S |
| **171Yb** | LYVE1 | EPR21857, Rabbit monoclonal | Sinusoidal endothelial cells and subsets of macrophages | Abcam/ab232935 |
| **176Yb** | EGFR | EP38Y, Rabbit monoclonal | Cell membrane | Abcam/ ab272293 |
| **176Yb** | Na/K ATPase | D4Y7E, Rabbit IgG | Cell membrane | Cell Signaling/ 23565S |
| **191 Ir** | Nuclear DNA | | Nuclei | Fluidigm/ 201192B |
| **196Pt** | Collagen I | EPR7785, Rabbit monoclonal | Connective tissue around portal triad | Abcam/ab215969 |

Table S3 Negative ion species identified by MALDI Orbitrap

| m/z | Species | Formula | Δppm | Adduct |
|---|---|---|---|---|
| 409.2354 | PA(16:0) | C19H39O7P | 1.6 | [M-H]- |
| 419.2561 | CPA(18:0) | C21H41O6P | 1.6 | [M-H]- |
| 427.2091 | PG(12:0) | C18H37O9P | 2.6 | [M-H]- |
| 435.251 | PA(18:1) | C21H41O7P | 1.6 | [M-H]- |
| 437.2667 | PA(18:0) | C21H43O7P | 1.6 | [M-H]- |
| 441.2248 | PG(13:0) | C19H39O9P | 2.4 | [M-H]- |
| 452.2776 | PE(16:0) | C21H44NO7P | 1.6 | [M-H]- |
| 455.2404 | PG(14:0) | C20H41O9P | 2.5 | [M-H]- |
| 464.3139 | PE(P-18:0) | C23H48NO6P | 1.6 | [M-H]- |
| 469.2559 | PG(15:0) | C21H43O9P | 2.8 | [M-H]- |
| 478.2931 | PE(18:1) | C23H46NO7P | 1.6 | [M-H]- |
| 480.3088 | PE(18:0) | C23H48NO7P | 1.6 | [M-H]- |
| 500.2783 | PE(20:4) | C25H44NO7P | 0 | [M-H]- |
| 504.3104 | PE(20:2) | C25H48NO7P | 1.7 | [M-H]- |
| 506.3261 | PE(20:1) | C25H50NO7P | 1.7 | [M-H]- |
| 508.3053 | PC(16:0) | C24H48NO8P | 1.7 | [M-H]- |
| 508.3408 | PE(20:0) | C25H52NO7P | 0.2 | [M-H]- |
| 524.2788 | PE(22:6) | C27H44NO7P | 1.1 | [M-H]- |
| 524.3 | PS(18:0) | C24H48NO9P | 1.1 | [M-H]- |
| 535.3026 | PG(20:2) | C26H49O9P | 2.9 | [M-H]- |
| 550.3135 | PS(20:1) | C26H50NO9P | 2.7 | [M-H]- |
| 552.5004 | Cer(d34:1(2OH)) | C34H67NO4 | 1.3 | [M-H]- |
| 553.3525 | PG(21:0) | C27H55O9P | 2.5 | [M-H]- |
| 561.3566 | PA(26:1) | C29H55O8P | 0.8 | [M-H]- |
| 571.2891 | PI(16:0) | C25H49O12P | 0.4 | [M-H]- |
| 579.3309 | PA-PA | C28H53O10P | 1 | [M-H]- |
| 581.3095 | POV-PG | C27H51O11P | 0.3 | [M-H]- |
| 583.3248 | PI(P-18:0) | C27H53O11P | 0.8 | [M-H]- |
| 587.3703 | PA(28:2) | C31H57O8P | 2.6 | [M-H]- |
| 594.3412 | PS-PC | C28H54NO10P | 0 | [M-H]- |
| 597.3047 | PI(18:1) | C27H51O12P | 0.2 | [M-H]- |
| 599.3196 | PI(18:0) | C27H53O12P | 1 | [M-H]- |
| 601.3872 | PA(29:2) | C32H59O8P | 0.5 | [M-H]- |
| 603.2941 | OKODiA-PA | C29H49O11P | 0.3 | [M-H]- |
| 605.3469 | OA-PA | C30H55O10P | 1.5 | [M-H]- |
| 616.4702 | CerP(d34:1) | C34H68NO6P | 1.6 | [M-H]- |
| 619.2884 | PI(20:4(5Z,8Z,11Z,14Z)/0:0) | C29H49O12P | 0.8 | [M-H]- |
| 622.3725 | PS(24:0) | C30H58NO10P | 0.2 | [M-H]- |
| 627.403 | PA(41:3) | C34H61O8P | 0.2 | [M-H]- |
| 634.3724 | OG-PC | C31H58NO10P | 0.3 | [M-H]- |
| 634.6524 | Cer(m42:0) | C42H85NO2 | 2.6 | [M-H]- |
| 644.5016 | CerP(d36:1) | C36H72NO6P | 1.4 | [M-H]- |
| 647.4657 | PA(16:0/16:0) | C35H69O8P | 0 | [M-H]- |
| 650.4037 | PS(26:0) | C32H62NO10P | 0.2 | [M-H]- |
| 664.4194 | PC(25:0(COOH)) | C33H64NO10P | 0.2 | [M-H]- |
| 669.45 | PA(34:3) | C37H67O8P | 0.1 | [M-H]- |
| 671.4651 | PA(16:0/18:2) | C37H69O8P | 1 | [M-H]- |
| 672.5328 | CerP(d38:1) | C38H76NO6P | 1.3 | [M-H]- |

| | | | | |
|---|---|---|---|---|
| 673.481 | PA(16:0/18:1) | C37H71O8P | 0.6 | [M-H]- |
| 674.4036 | PS(28:2) | C34H62NO10P | 0.3 | [M-H]- |
| 685.4812 | PA(25:2) | C38H71O8P | 0.3 | [M-H]- |
| 687.4971 | PA(25:1) | C38H73O8P | 0.1 | [M-H]- |
| 687.544 | SM(d18:1/16:0) | C38H77N2O6P | 0.9 | [M-CH3]- |
| 690.435 | PS(29:1) | C35H66NO10P | 0.3 | [M-H]- |
| 693.4505 | PA(36:5) | C39H67O8P | 0.6 | [M-H]- |
| 695.4651 | PA(16:0/20:4) | C39H69O8P | 1 | [M-H]- |
| 697.4808 | PA(36:3) | C39H71O8P | 0.8 | [M-H]- |
| 699.4965 | PA(18:1/18:1) | C39H73O8P | 0.8 | [M-H]- |
| 700.5645 | CerP(d40:1) | C40H80NO6P | 0.8 | [M-H]- |
| 701.5119 | PA(18:0/18:1) | C39H75O8P | 1.1 | [M-H]- |
| 709.4813 | PA(37:4) | C40H71O8P | 0.1 | [M-H]- |
| 712.492 | PE(34:3) | C39H72NO8P | 0.4 | [M-H]- |
| 713.5125 | PA(27:2) | C40H75O8P | 0.2 | [M-H]- |
| 714.5071 | PE(34:2) | C39H74NO8P | 1.2 | [M-H]- |
| 715.4917 | PG(P-33:2) | C39H73O9P | 0.3 | [M-H]- |
| 715.5748 | SM(d36:1) | C40H81N2O6P | 1.6 | [M-CH3]- |
| 716.5233 | PE(16:0/18:1) | C39H76NO8P | 0.5 | [M-H]- |
| 718.5387 | PE(16:0/18:0) | C39H78NO8P | 0.8 | [M-H]- |
| 719.465 | PA(38:6) | C41H69O8P | 1 | [M-H]- |
| 721.4809 | PA(38:5) | C41H71O8P | 0.6 | [M-H]- |
| 723.4964 | PA(18:0/20:4) | C41H73O8P | 0.9 | [M-H]- |
| 725.5124 | PA(38:3) | C41H75O8P | 0.4 | [M-H]- |
| 726.58 | CerP(d42:2) | C42H82NO6P | 0.9 | [M-H]- |
| 727.4551 | PG(33:4) | C39H69O10P | 0.7 | [M-H]- |
| 728.5232 | PC(32:2) | C40H76NO8P | 0.5 | [M-H]- |
| 728.5958 | CerP(d32:1) | C42H84NO6P | 0.7 | [M-H]- |
| 731.4852 | PG(33:2) | C39H73O10P | 2.3 | [M-H]- |
| 733.4814 | PA(29:6) | C42H71O8P | 0.1 | [M-H]- |
| 736.492 | PE(26:5) | C41H72NO8P | 0.4 | [M-H]- |
| 738.5071 | PE(16:0/20:4) | C41H74NO8P | 1.1 | [M-H]- |
| 740.5232 | PE(16:0/20:3)/PE(18:1/18:2) | C41H76NO8P | 0.6 | [M-H]- |
| 742.5385 | PE(18:0/18:2) | C41H78NO8P | 1 | [M-H]- |
| 743.6062 | SM(d38:1) | C42H85N2O6P | 1.4 | [M-CH3]- |
| 744.5551 | PE(36:1) | C41H80NO8P | 0.3 | [M-H]- |
| 745.481 | PA(40:6) | C43H71O8P | 0.5 | [M-H]- |
| 745.5023 | PG(44:2) | C40H75O10P | 0.2 | [M-H]- |
| 746.5712 | PE(36:0) | C41H82NO8P | 0.8 | [M-H]- |
| 746.7046 | Cer(d34:1) | C48H93NO4 | 1.9 | [M-H]- |
| 747.4964 | PA(18:0/22:6) | C43H73O8P | 0.9 | [M-H]- |
| 747.5176 | PG(44:1) | C40H77O10P | 0.7 | [M-H]- |
| 749.5126 | PA(40:5) | C43H75O8P | 0.2 | [M-H]- |
| 750.5432 | PE(P-18:0/20:4) | C43H78NO7P | 1.5 | [M-H]- |
| 752.5232 | PC(34:4) | C42H76NO8P | 0.5 | [M-H]- |
| 752.5601 | PE(O-38:4) | C43H80NO7P | 0.2 | [M-H]- |
| 756.5182 | PS(P-18:0/17:2(9Z,12Z)) | C41H76NO9P | 0.4 | [M-H]- |
| 756.5545 | PC(34:2) | C42H80NO8P | 0.5 | [M-H]- |
| 758.5338 | PS(P-25:1) | C41H78NO9P | 0.5 | [M-H]- |
| 760.4919 | PE(38:7) | C43H72NO8P | 0.5 | [M-H]- |

| | | | | |
|---|---|---|---|---|
| 762.507 | PE(16:0/22:6) | C43H74NO8P | 1.2 | [M-H]- |
| 764.5234 | PE(38:5) | C43H76NO8P | 0.2 | [M-H]- |
| 766.5384 | PE(18:0/20:4) | C43H78NO8P | 1.1 | [M-H]- |
| 768.5548 | PE(18:0/20:3) | C43H80NO8P | 0.1 | [M-H]- |
| 769.6217 | SM(d42:2) | C44H87N2O6P | 1.5 | [M-H]- |
| 770.534 | PS(P-36:2) | C42H78NO9P | 0.1 | [M-H]- |
| 770.5698 | PE(18:0/18:2) | C43H82NO8P | 0.9 | [M-H]- |
| 771.517 | PG(36:3) | C42H77O10P | 1.4 | [M-H]- |
| 771.6376 | SM(d18:1/22:0) | C44H89N2O6P | 1.2 | [M-CH3]- |
| 773.5129 | PA(42:7) | C45H75O8P | 0.3 | [M-H]- |
| 773.533 | PG(18:1/18:1) | C42H79O10P | 1.1 | [M-H]- |
| 777.5646 | PG(36:0) | C42H83O10P | 0.7 | [M-H]- |
| 778.5024 | PE(38:6(14OH)) | C43H74NO9P | 0.5 | [M-H]- |
| 780.5181 | PE(38:4(15Ke)) | C43H76NO9P | 0.5 | [M-H]- |
| 780.5544 | PC(36:4) | C44H80NO8P | 0.6 | [M-H]- |
| 782.4971 | PS(36:4) | C42H74NO10P | 0.8 | [M-H]- |
| 782.5337 | PE(38:4(12OH[S])) | C43H78NO9P | 0.6 | [M-H]- |
| 784.5494 | PS(P-37:2) | C43H80NO9P | 0.4 | [M-H]- |
| 785.6534 | SM(d40:1) | C45H91N2O6P | 1.1 | [M-H]- |
| 786.5077 | PE(40:8) | C45H74NO8P | 0.3 | [M-H]- |
| 786.5285 | PS(18:0/18:2) | C42H78NO10P | 0.7 | [M-H]- |
| 788.523 | PE(40:7) | C45H76NO8P | 0.8 | [M-H]- |
| 788.5439 | PS(18:0/18:1) | C42H80NO10P | 1 | [M-H]- |
| 790.5027 | PS(P-38:6) | C44H74NO9P | 0.2 | [M-H]- |
| 790.5384 | PE(18:0/22:6) | C45H78NO8P | 1.1 | [M-H]- |
| 792.554 | PE(18:1/22:4) | C45H80NO8P | 1.1 | [M-H]- |
| 794.5329 | PS(P-38:4) | C44H78NO9P | 1.6 | [M-H]- |
| 794.5698 | PE(18:0/22:4) | C45H82NO8P | 0.9 | [M-H]- |
| 796.5857 | PE(40:3) | C45H84NO8P | 0.6 | [M-H]- |
| 797.6533 | SM(d42:2) | C46H91N2O6P | 1.1 | [M-CH3]- |
| 799.669 | SM(d42:1) | C46H93N2O6P | 1 | [M-CH3]- |
| 802.5599 | PT(36:1) | C43H82NO10P | 0.6 | [M-H]- |
| 804.5546 | PC(38:6) | C46H80NO8P | 0.3 | [M-H]- |
| 806.4971 | PS(38:6) | C44H74NO10P | 0.8 | [M-H]- |
| 806.5336 | PE(40:6(14OH)) | C45H78NO9P | 0.6 | [M-H]- |
| 810.5281 | PS(18:0/20:4) | C44H78NO10P | 1.2 | [M-H]- |
| 814.5388 | PE(42:8) | C47H78NO8P | 0.5 | [M-H]- |
| 816.5545 | PE(42:7) | C47H80NO8P | 0.5 | [M-H]- |
| 818.533 | PS(P-40:6) | C46H78NO9P | 1.4 | [M-H]- |
| 818.5701 | PE(42:6) | C47H82NO8P | 0.6 | [M-H]- |
| 833.5175 | PI(16:0/18:2) | C43H79O13P | 1.2 | [M-H]- |
| 834.5276 | PS(18:0/22:6) | C46H78NO10P | 1.8 | [M-H]- |
| 835.5323 | PI(44:1) | C43H81O13P | 2.2 | [M-H]- |
| 857.5176 | PI(16:0/20:4) | C45H79O13P | 1.1 | [M-H]- |
| 859.5335 | PI(36:3/22:2) | C45H81O13P | 0.8 | [M-H]- |
| 861.5489 | PI(18:0/18:2) | C45H83O13P | 1.1 | [M-H]- |
| 863.5638 | PI(26:1) | C45H85O13P | 1.9 | [M-H]- |
| 871.5334 | PI(27:4) | C46H81O13P | 1 | [M-H]- |
| 872.638 | PS(42:1) | C48H92NO10P | 0.7 | [M-H]- |
| 881.5181 | PI(38:6) | C47H79O13P | 0.6 | [M-H]- |

| | | | | |
|---|---|---|---|---|
| 883.5333 | PI(18:1/20:4) | C47H81O13P | 1 | [M-H]- |
| 885.5482 | PI(18:0/20:4) | C47H83O13P | 1.9 | [M-H]- |
| 899.5648 | PI(39:4) | C48H85O13P | 0.8 | [M-H]- |
| 909.5488 | PI(18:0/22:6) | C49H83O13P | 1.1 | [M-H]- |
| 911.5647 | PI(18:0/22:5) | C49H85O13P | 0.8 | [M-H]- |
| 913.5806 | PI(40:4) | C49H87O13P | 0.6 | [M-H]- |
| 1050.526 | CDP-DG(40:7) | C52H83N3O15P2 | 3.4 | [M-H]- |
| 1052.54 | CDP-DG(40:6) | C52H85N3O15P2 | 1.8 | [M-H]- |
| 1447.966 | CL(1'-[18:2/18:2/18:2/18:2]) | C81H142O17P2 | 0.6 | [M-H]- |
| 1454.875 | GalNAcbeta1-4(NeuGcalpha2-3)Galbeta1-4Glcbeta-Cer(d40:1) | C71H129N3O27 | 0.7 | [M-H]- |
| 1482.906 | GalNAcbeta1-4(NeuGcalpha2-3)Galbeta1-4Glcbeta-Cer(d42:1) | C73H133N3O27 | 0.4 | [M-H]- |

Table S4 Positive ion species identified by MALDI Orbitrap

| m/z | Species | Formula | Δppm | Adduct |
|---|---|---|---|---|
| 352.2249 | C16 Sphingosine-1-phosphate | C16H34NO5P | 0.5 | [M+H]+ |
| 367.336 | Cholesterol species | C27H44O | 0.2 | [M+H-H2O]+ |
| 369.3516 | Cholesterol species | C27H46O | 0 | [M+H-H2O]+ |
| 383.3309 | Cholesterol species | C27H44O2 | 0.2 | [M+H-H2O]+ |
| 385.3466 | Cholesterol species | C27H46O2 | 0.2 | [M-H]+ |
| 401.3414 | Cholesterol species | C27H46O3 | 0.1 | [M+H-H2O]+ |
| 469.1659 | Ptilosteroid B | C21H34O7S | 0.5 | [M+K]+ |
| 469.1758 | PA(18:4(6Z,9Z,12Z,15Z)/0:0) | C21H35O7P | 1.4 | [M+K]+ |
| 478.3292 | PE(P-19:1(12Z)/0:0) | C24H48NO6P | 0.1 | [M+H]+ |
| 496.3398 | PC(16:0) | C24H50NO7P | 0 | [M+H]+ |
| 506.3605 | PC(P-18:1) | C26H52NO6P | 0.1 | [M+H]+ |
| 516.2851 | PE(P-19:1) | C24H48NO6P | 0.1 | [M+K]+ |
| 518.3218 | PC(16:0) | C24H50NO7P | 0.1 | [M+Na]+ |
| 520.5088 | Cer(d34:1) | C34H67NO3 | 0.1 | [M+H-H2O]+ |
| 522.3554 | PC(18:1) | C26H52NO7P | 0 | [M+H]+ |
| 523.2643 | Vitamine D3 species | C27H36O3F6 | 0.3 | [M+H]+ |
| 524.371 | PC(18:0) | C26H54NO7P | 0.1 | [M+H]+ |
| 534.2956 | PC(0:0/16:0) | C24H50NO7P | 0.1 | [M+K]+ |
| 546.3531 | PC(18:0) | C26H54NO7P | 0.2 | [M+Na]+ |
| 550.3503 | PC(19:1) | C27H52NO8P | 0 | [M+H]+ |
| 562.3269 | PC(18:0) | C26H54NO7P | 0.1 | [M+K]+ |
| 580.3609 | PS(22:1) | C28H54NO9P | 0 | [M+H]+ |
| 588.3062 | PC(19:1) | C27H52NO8P | 0 | [M+K]+ |
| 594.3765 | PC(21:0(CHO)) | C29H56NO9P | 0 | [M+H]+ |
| 604.6027 | Cer(d40:1) | C40H79NO3 | 0 | [M+H-H2O]+ |
| 608.3922 | PON-PE | C30H58NO9P | 0.1 | [M+H]+ |
| 618.24 | Taurocholic acid 3-sulfate | C26H45NO10S2 | 3.7 | [M+Na]+ |
| 618.6184 | Cer(d41:1) | C41H81NO3 | 0 | [M+H-H2O]+ |
| 622.4443 | PC(24:0) | C32H64NO8P | 0 | [M+H]+ |
| 630.6184 | Cer(d42:2) | C42H81NO3 | 0 | [M+H-H2O]+ |
| 632.3324 | PC(21:0(CHO)) | C29H56NO9P | 0 | [M+K]+ |
| 632.6341 | Cer(d42:1) | C42H83NO3 | 0.1 | [M+H-H2O]+ |
| 639.2138 | Ptilosaponoside A | C27H44O14S2 | 0.2 | [M+H-H2O]+ |
| 650.4391 | PON-PC | C33H64NO9P | 0.1 | [M+H]+ |

| | | | | |
|---|---|---|---|---|
| 655.2295 | Halistanol sulfonic acid A | C28H48O12S3 | 3 | [M+H-H2O]+ |
| 666.434 | PC(25:0(COOH)) | C33H64NO10P | 0 | [M+H]+ |
| 672.4213 | PON-PC | C33H64NO9P | 0.2 | [M+Na]+ |
| 673.5895 | 18:1 Cholesterol ester | C45H78O2 | 0.2 | [M+Na]+ |
| 675.5435 | SM(d32:1) | C37H75N2O6P | 0 | [M+H]+ |
| 678.4704 | PS(P-29:0) | C35H68NO9P | 0 | [M+H]+ |
| 687.5475 | 18:2 Cholesterol ester | C45H76O2 | 0.3 | [M+K]+ |
| 688.395 | PON-PC | C33H64NO9P | 0.1 | [M+K]+ |
| 688.4162 | PC(25:0(COOH)) | C33H64NO10P | 0.2 | [M+Na]+ |
| 689.5606 | SM(d16:1/17:0) | C38H77N2O6P | 2.1 | [M+H]+ |
| 701.5592 | SM(d16:1/18:1) | C39H77N2O6P | 0 | [M+H]+ |
| 703.5748 | SM(d34:1) | C39H79N2O6P | 0.1 | [M+H]+ |
| 704.3899 | PC(25:0(COOH)) | C33H64NO10P | 0.1 | [M+K]+ |
| 716.4264 | PS(P-29:0) | C35H68NO9P | 0 | [M+K]+ |
| 716.5224 | PE(34:2) | C39H74NO8P | 0.1 | [M+H]+ |
| 717.5906 | SM(d18:1/17:0) | C40H81N2O6P | 0.1 | [M+H]+ |
| 718.5382 | PE(34:1) | C39H76NO8P | 0.1 | [M+H]+ |
| 721.4779 | PA(36:3) | C39H71O8P | 0 | [M+Na]+ |
| 723.4935 | PA(36:2) | C39H73O8P | 0 | [M+Na]+ |
| 725.5567 | SM(d34:1) | C39H79N2O6P | 0.1 | [M+Na]+ |
| 731.6061 | SM(d36:1) | C41H83N2O6P | 0 | [M+H]+ |
| 732.5537 | PC(32:1) | C40H78NO8P | 0 | [M+H]+ |
| 734.5695 | PC(32:0) | C40H80NO8P | 0.1 | [M+H]+ |
| 737.4517 | PA(36:3) | C39H71O8P | 0.1 | [M+K]+ |
| 739.4675 | PA(36:2) | C39H73O8P | 0.1 | [M+K]+ |
| 740.5221 | PE(36:4) | C41H74NO8P | 0.5 | [M+H]+ |
| 741.5302 | SM(d34:1) | C39H79N2O6P | 0.7 | [M+K]+ |
| 742.5389 | PE(36:3) | C41H76NO8P | 2.5 | [M+H]+ |
| 744.5538 | PE(36:2) | C41H78NO8P | 0 | [M+H]+ |
| 746.5697 | PE(36:1) | C41H80NO8P | 0.4 | [M+H]+ |
| 753.5881 | SM(d36:1) | C41H83N2O6P | 0 | [M+Na]+ |
| 754.4784 | PE(34:2) | C39H74NO8P | 0 | [M+K]+ |
| 754.5365 | PC(32:1) | C40H78NO8P | 1 | [M+Na]+ |
| 756.5529 | PC(34:3) | C42H78NO8P | 1.2 | [M+H]+ |
| 758.5693 | PC(34:2) | C42H80NO8P | 0.1 | [M+H]+ |
| 759.6374 | SM(d38:1) | C43H87N2O6P | 0 | [M+H]+ |
| 760.5854 | PC(34:1) | C42H82NO8P | 0.5 | [M+H]+ |
| 761.4515 | PA(38:5) | C41H71O8P | 0.5 | [M+K]+ |
| 763.4674 | PA(38:4) | C41H73O8P | 0.1 | [M+K]+ |
| 764.5222 | PE(38:6) | C43H74NO8P | 0.4 | [M+H]+ |
| 765.483 | PA(38:3) | C41H75O8P | 0.2 | [M+K]+ |
| 766.5375 | PE(38:5) | C43H76NO8P | 0.8 | [M+H]+ |
| 766.5516 | CerP(d42:2) | C42H82NO6P | 0.6 | [M+K]+ |
| 768.5537 | PE(38:4) | C43H78NO8P | 0.1 | [M+H]+ |
| 770.5097 | PC(32:1) | C40H78NO8P | 0 | [M+K]+ |
| 770.5681 | PE(36:0) | C41H82NO8P | 1.3 | [M+Na]+ |
| 772.5253 | PC(32:0) | C40H80NO8P | 0 | [M+K]+ |
| 772.5851 | PE(38:2) | C43H82NO8P | 0 | [M+H]+ |
| 773.6531 | SM(d39:1) | C44H89N2O6P | 0 | [M+H]+ |
| 774.5643 | PS(O-36:2) | C42H80NO9P | 0 | [M+H]+ |

| | | | | |
|---|---|---|---|---|
| 774.6008 | PE(38:1) | C43H84NO8P | 0.1 | [M+H]+ |
| 776.5799 | PS(O-36:1) | C42H82NO9P | 0.1 | [M+H]+ |
| 778.4782 | PE(36:4) | C41H74NO8P | 0.2 | [M+K]+ |
| 778.5362 | PC(34:3) | C42H78NO8P | 0.6 | [M+Na]+ |
| 780.5515 | PC(34:2) | C42H80NO8P | 0.2 | [M+Na]+ |
| 781.6194 | SM(d38:1) | C43H87N2O6P | 0 | [M+Na]+ |
| 782.5097 | PE(36:2) | C41H78NO8P | 0.1 | [M+K]+ |
| 782.5687 | PC(36:4) | C44H80NO8P | 0.9 | [M+H]+ |
| 784.5254 | PE(36:1) | C41H80NO8P | 0.1 | [M+K]+ |
| 784.5858 | PC(36:3) | C44H82NO8P | 1 | [M+H]+ |
| 785.6531 | SM(d40:2) | C45H89N2O6P | 0 | [M+H]+ |
| 786.6008 | PE(36:2) | C44H84NO8P | 0.1 | [M+H]+ |
| 787.6687 | SM(d40:1) | C45H91N2O6P | 0.1 | [M+H]+ |
| 788.6173 | PC(36:1) | C44H86NO8P | 1.2 | [M+H]+ |
| 790.5362 | PE(38:4) | C43H78NO8P | 0.6 | [M+Na]+ |
| 790.5593 | PS(36:1) | C42H80NO10P | 0 | [M+H]+ |
| 792.5534 | PE(40:6) | C45H78NO8P | 0.5 | [M+H]+ |
| 794.5097 | PC(34:3) | C42H78NO8P | 0.1 | [M+K]+ |
| 794.5685 | PE(40:5) | C45H80NO8P | 1.2 | [M+H]+ |
| 794.5828 | CerP(d44:2) | C44H86NO6P | 0.5 | [M+K]+ |
| 796.5252 | PC(34:2) | C42H80NO8P | 0.1 | [M+K]+ |
| 796.5845 | PE(40:4) | C45H82NO8P | 0.7 | [M+H]+ |
| 797.592 | SM(d38:1) | C43H87N2O6P | 1.7 | [M+K]+ |
| 798.5412 | PC(16:1/18:0) | C42H82NO8P | 0.3 | [M+K]+ |
| 801.6844 | SM(d41:1) | C46H93N2O6P | 0 | [M+H]+ |
| 802.4782 | PE(38:6) | C43H74NO8P | 0.2 | [M+K]+ |
| 804.4933 | PE(38:5) | C43H76NO8P | 0.9 | [M+K]+ |
| 804.5514 | PC(36:4) | C44H80NO8P | 0.1 | [M+Na]+ |
| 806.5096 | PE(38:4) | C43H78NO8P | 0.1 | [M+K]+ |
| 806.5686 | PC(36:3) | C44H82NO8P | 1.9 | [M+Na]+ |
| 808.5696 | GlcCer(d38:2(2OH)) | C44H83NO9 | 0.4 | [M+K]+ |
| 808.5839 | PC(36:2) | C44H84NO8P | 1.5 | [M+Na]+ |
| 809.6507 | SM(d40:1) | C45H91N2O6P | 0.1 | [M+Na]+ |
| 810.5411 | PE(38:2) | C43H82NO8P | 0.2 | [M+K]+ |
| 810.6007 | PC(38:4) | C46H84NO8P | 0 | [M+H]+ |
| 812.52 | PE(40:7) | C45H76NO8P | 0.1 | [M+Na]+ |
| 812.5421 | PS(36:1) | C42H80NO10P | 1.1 | [M+Na]+ |
| 812.6163 | PC(38:3) | C46H86NO8P | 0.1 | [M+H]+ |
| 813.6844 | SM(d42:2) | C47H93N2O6P | 0.1 | [M+H]+ |
| 814.5588 | PS(38:3) | C44H80NO10P | 0.6 | [M+H]+ |
| 815.7002 | SM(d42:1) | C47H95N2O6P | 0.1 | [M+H]+ |
| 818.5097 | PC(36:5) | C44H78NO8P | 0 | [M+K]+ |
| 820.5251 | PC(36:4) | C44H80NO8P | 0.2 | [M+K]+ |
| 822.5413 | PC(36:3) | C44H82NO8P | 0.4 | [M+K]+ |
| 823.6663 | SM(d41:1) | C46H93N2O6P | 0.1 | [M+Na]+ |
| 824.5566 | PC(36:2) | C44H84NO8P | 0 | [M+K]+ |
| 825.6244 | SM(d40:1) | C45H91N2O6P | 0.2 | [M+K]+ |
| 826.5723 | PC(36:1) | C44H86NO8P | 0 | [M+K]+ |
| 828.5151 | PS(14:0/22:1(11Z)) | C42H80NO10P | 0.1 | [M+K]+ |
| 828.5515 | PC(38:6) | C46H80NO8P | 0.1 | [M+Na]+ |

| | | | | |
|---|---|---|---|---|
| 830.5674 | PC(38:5) | C46H82NO8P | 0.5 | [M+Na]+ |
| 832.5831 | PC(38:4) | C46H84NO8P | 0.6 | [M+Na]+ |
| 834.6003 | PC(40:6) | C48H84NO8P | 0.5 | [M+H]+ |
| 835.6663 | SM(d42:2) | C47H93N2O6P | 0 | [M+Na]+ |
| 837.682 | SM(d42:1) | C47H95N2O6P | 0 | [M+Na]+ |
| 839.64 | SM(d41:1) | C46H93N2O6P | 0.4 | [M+K]+ |
| 840.6436 | PC(38:0) | C46H92NO8P | 2 | [M+Na]+ |
| 844.5252 | PC(38:6) | C46H80NO8P | 0.1 | [M+K]+ |
| 846.5412 | PC(38:5) | C46H82NO8P | 0.2 | [M+K]+ |
| 848.5565 | PC(38:4) | C46H84NO8P | 0.2 | [M+K]+ |
| 850.5721 | PC(38:3) | C46H86NO8P | 0.2 | [M+K]+ |
| 851.6402 | SM(d42:2) | C47H93N2O6P | 0.1 | [M+K]+ |
| 853.6559 | SM(d42:1) | C47H95N2O6P | 0.1 | [M+K]+ |
| 869.6994 | TG(50:2) | C53H98O6 | 0.1 | [M+K]+ |
| 872.5569 | PC(40:6) | C48H84NO8P | 0.4 | [M+K]+ |
| 879.7412 | TG(52:3) | C55H100O6 | 0 | [M+Na]+ |
| 881.7568 | TG(52:2) | C55H102O6 | 0.1 | [M+Na]+ |
| 893.6994 | TG(52:4) | C55H98O6 | 0.1 | [M+K]+ |
| 895.7151 | TG(52:3) | C55H100O6 | 0.1 | [M+K]+ |
| 897.7306 | TG(52:2) | C55H102O6 | 0.2 | [M+K]+ |
| 919.7151 | TG(54:5) | C57H100O6 | 0.1 | [M+K]+ |
| 921.7306 | TG(54:4) | C57H102O6 | 0.2 | [M+K]+ |
| 923.7462 | TG(54:3) | C57H104O6 | 0.2 | [M+K]+ |
| 925.5202 | PI(38:4) | C47H83O13P | 0.1 | [M+K]+ |

MSMS in human liver
MSMS in mouse liver

Supplementary Videos

**Video S1 3D reconstruction of human liver tissue section.** Selected lipid species are overlaid and pseudocolored for visualization. Magenta PC32:1; red SM 42:2; green PC 32:0; yellow TG 52:2.

**Video S2 3D reconstruction of human liver portal triad.** The selected metabolites and lipids are fused with protein markers on the same serial tissues. Cyan, CD4; Red, CD45; Orange, CD68; Blue, SMA; Lime green, GSH; Green, 7-[(1S, 2S)-2-(Heptylamine) cyclohexyl] heptanoic acid tauodexycholic acid; Yellow, HemeB; Meganta, PS 36:1.
